# RhoBAST - a rhodamine-binding aptamer for super-resolution RNA imaging

**DOI:** 10.1101/2020.03.12.988782

**Authors:** Murat Sunbul, Jens Lackner, Annabell Martin, Daniel Englert, Benjamin Hacene, Karin Nienhaus, G. Ulrich Nienhaus, Andres Jäschke

**Affiliations:** Institute of Pharmacy and Molecular Biotechnology (IPMB), Heidelberg University, D-69120 Heidelberg, Germany; Institute of Applied Physics (APH), Karlsruhe Institute of Technology (KIT), Wolfgang-Gaede-Str. 1, D-76131 Karlsruhe, Germany; Institute of Nanotechnology (INT), Karlsruhe Institute of Technology (KIT), D-76344 Eggenstein-Leopoldshafen, Germany; Institute of Biological and Chemical Systems (IBCS), Karlsruhe Institute of Technology (KIT), D-76344 Eggenstein-Leopoldshafen, Germany; Department of Physics, University of Illinois at Urbana−Champaign, 1110 West Green Street, Urbana, Illinois 61801, United States

## Abstract

RhoBAST is a novel fluorescence light-up RNA aptamer (FLAP) that transiently binds a fluorogenic rhodamine dye. Fast dye association and dissociation result in intermittent fluorescence emission, facilitating single-molecule localization microscopy (SMLM) with an image resolution not limited by photobleaching. We demonstrate RhoBAST's excellent properties as a RNA marker by resolving subcellular and subnuclear structures of RNA in live and fixed cells by SMLM and structured illumination microscopy (SIM).

## Main

Super-resolution fluorescence microscopy of cellular structures continually provides fundamentally new insights into biology^1^. Over the years, an elaborate toolbox has become available to attach fluorescent markers to proteins for such studies. By contrast, marker tools for super-resolution RNA imaging are still scarce^2^. Recently, FLAPs have been recognized as promising tools for RNA imaging^3^. So far, they have mostly been employed for conventional, diffraction-limited imaging. FLAPs for super-resolution imaging have been reported only last year, specifically, the SiRA aptamer^4^ for stimulated emission depletion (STED) microscopy and the Pepper aptamer^5^ for structured illumination microscopy (SIM). Further optimization of these RNA markers will facilitate their widespread use in super-resolution microscopy.

Our laboratory has developed FLAPs that bind fluorophore-quencher conjugates and thereby disrupt contact quenching (Fig. 1a)^6, 7^. This strategy affords the use of bright and photostable state-of-the-art fluorophores for RNA imaging^7, 8^. For example, the SRB-2 aptamer binds various membrane-permeable rhodamine dyes with high affinity, yet it does not provide sufficient sensitivity for imaging RNAs of low abundance in cells^9^. Here we present RhoBAST, a FLAP endowed with excellent properties for single-molecule localization microscopy (SMLM)^10^, and demonstrate its ability to resolve subnuclear structures of RNA in live and fixed cells using SMLM and SIM.

**Figure 1.**
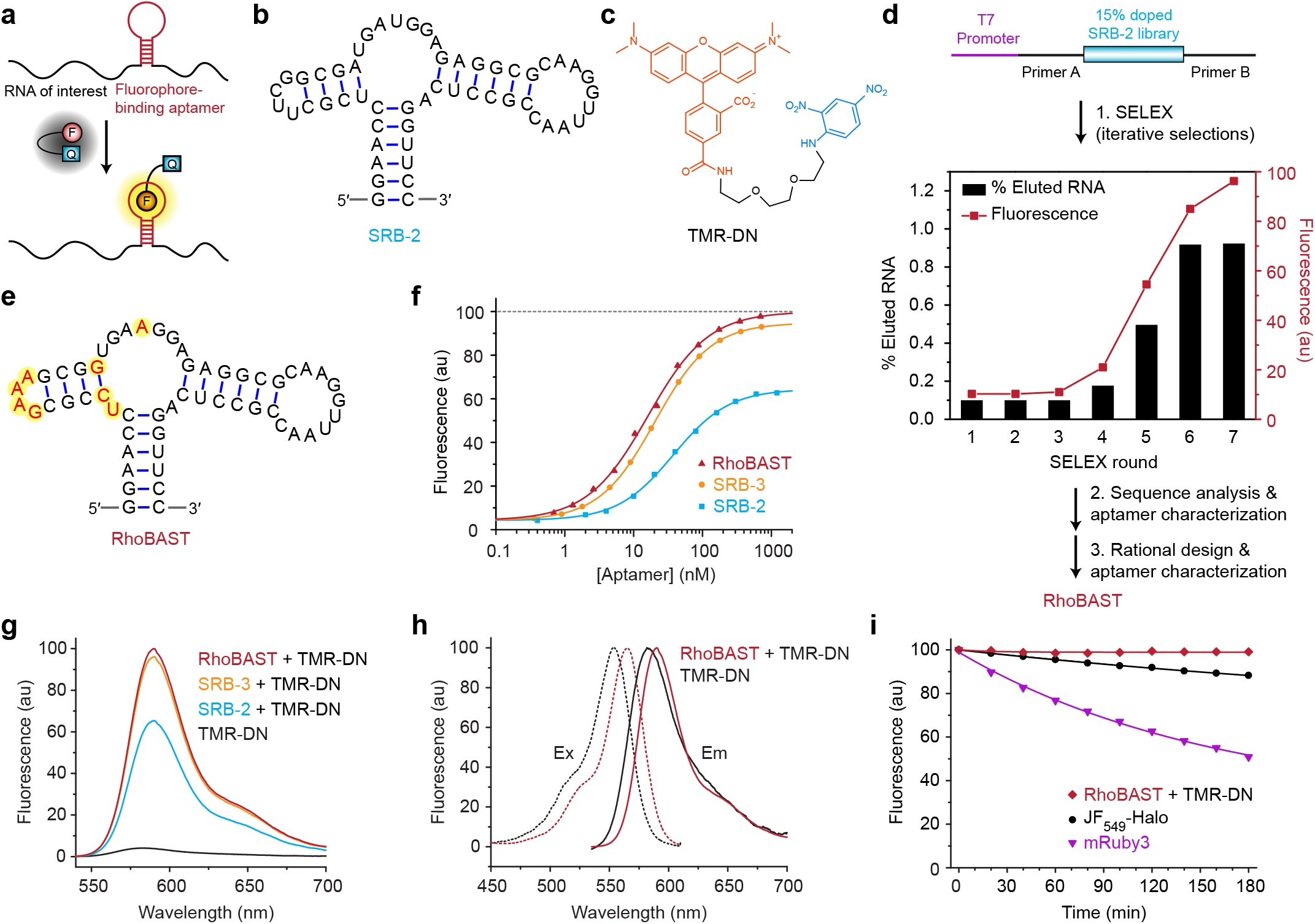
Directed evolution and characterization of RhoBAST. **a)** Scheme showing the use of a genetically encoded fluorophore-binding aptamer and a contact-quenched fluorophore-quencher conjugate (F-Q) for imaging an RNA of interest. F-Q is non-fluorescent in solution and lights up upon binding to the aptamer. **b)** Predicted secondary structure of SRB-2. **c)** Chemical structure of TMR-DN. **d)** Directed evolution of RhoBAST for binding to the TMR fluorophore using iterative cycles of SELEX and rational design. The initial library was composed of 15% doped SRB-2 flanked by constant primer A and primer B binding regions. The T7 promoter was used for efficient transcription of the library by T7 RNA polymerase. Progress of the selection was assessed by calculating the percentage of eluted RNA and the fluorescence enhancement of TMR-DN upon addition of RNA pools after each round. **e)** Predicted secondary structure of RhoBAST; changed ribonucleotides are highlighted in yellow. **f)** Binding isotherms resulting in equilibrium dissociation coefficients, *K*_D_, of 35 ± 1 nM for SRB-2, 20 ± 1 nM for SRB-3 and 15 ± 1 nM for RhoBAST, using TMR-DN as the ligand. **g)** Emission spectra of TMR-DN, SRB-2:TMR-DN, SRB-3:TMR-DN and RhoBAST:TMR-DN, all measured under identical conditions and normalized such that the RhoBAST:TMR-DN spectrum peaks at 100. **h)** Normalized excitation and emission spectra of TMR-DN and RhoBAST:TMR-DN. **i)** Fluorescence intensity decay of TMR-DN (20 nM) complexed with RhoBAST (500 nM), the JF_549_-Halo fluorophore (20 nM) and the mRuby3 fluorescent protein (20 nM). After 3 h, the fractional decrease was 0.9 ± 0.4%, 11.7 ± 0.9% and 49.5 ± 2.7% (mean ± s.d.), respectively.

Starting from a 15% doped SRB-2 aptamer library, *in vitro* evolution was employed to improve the affinity, thermal stability, photostability and brightness of SRB-2 using tetramethylrhodamine (TMR) as bait (Fig. 1b,c). Following incubation of the library with TMR-decorated beads, mutants binding TMR were specifically eluted, amplified and subsequently used as input for the next round of evolution. After seven iterations (Fig. 1d), the enriched variants were sequenced and screened for fluorescence enhancement upon binding the dye-quencher conjugate TMR-dinitroaniline (TMR-DN, Supplementary Fig. 1). Sequence and truncation analysis (Supplementary Figs. 2 and 3) revealed two beneficial mutations in the SRB-2 sequence, namely an insertion of a single U nucleotide between C6 and U7 and a U22A mutation (Supplementary Fig. 4). These features were incorporated into the design of SRB-3 (Supplementary Fig. 4), which exhibits 50% higher brightness and 1.8-fold higher affinity (*K*_D_ = 20 ± 1 nM) for TMR-DN than SRB-2 (Fig. 1f).

**Figure 2.**
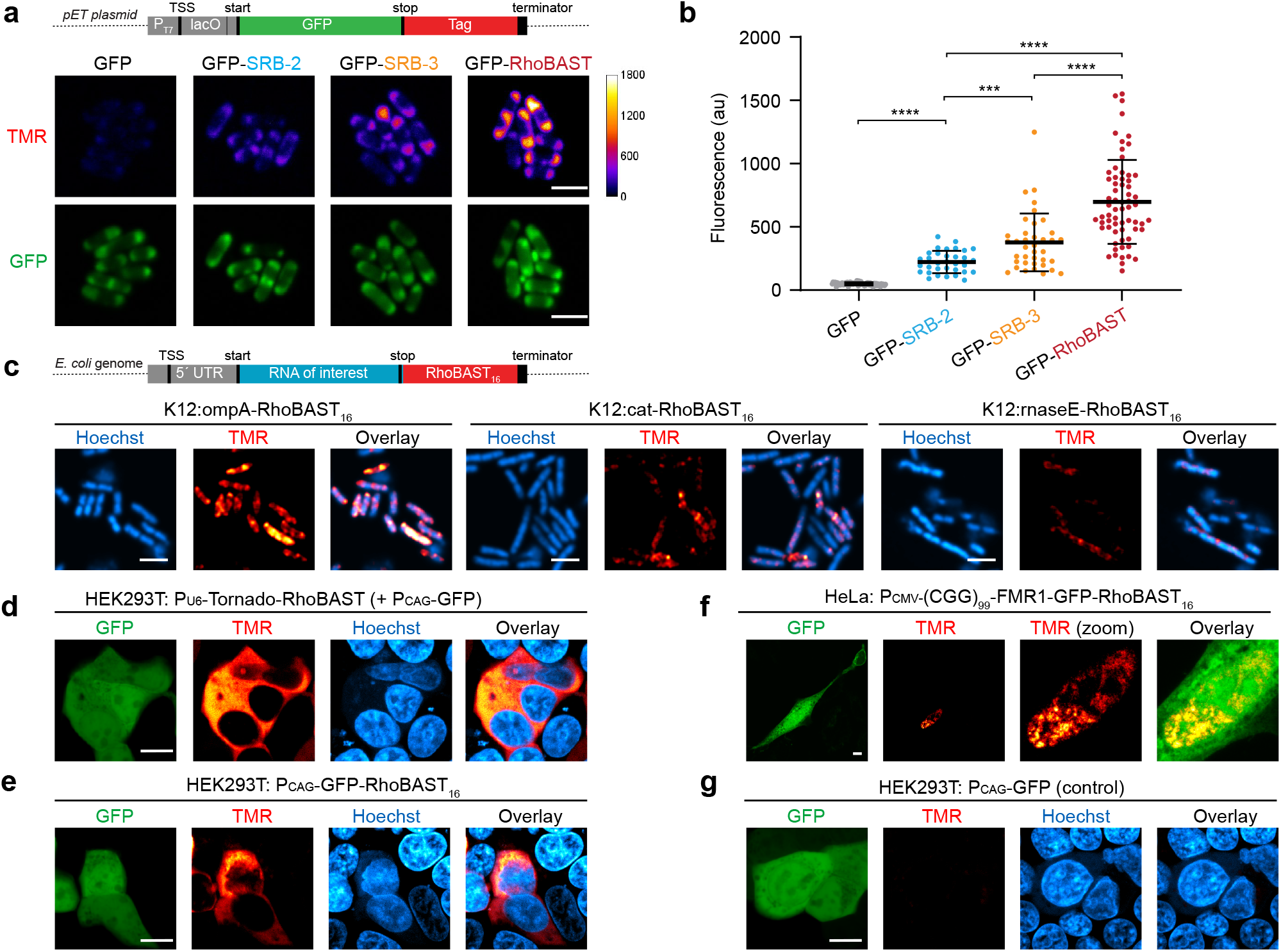
Performance of RhoBAST as a marker for RNA imaging in live cells. **a)** Live cell imaging of *gfp* mRNA (control) and *gfp* mRNA fused to a single copy of SRB-2, SRB-3 or RhoBAST. Bacteria were transformed with the pET plasmid carrying the *gfp* gene fused to the aptamer tags at the 3′ UTR. Scale bars, 3 μm. **b)** Quantification of the average TMR fluorescence at the poles of bacteria expressing the different RNA constructs. Each dot represents a single cell; also indicated are the means ± s.d. (N > 30 cells). Statistical comparison was performed by using a two-tailed t-test. ***, *P* ≤ 0.001; ****, *P* ≤ 0.0001. **c)** Live-cell mRNA imaging of endogenously RhoBAST-tagged *ompA*, *cat* and *rnaseE* in *E. coli* K12. **d)** Live-cell imaging of circular RhoBAST aptamers transcribed under the control of the U6 promoter using the Tornado system in HEK293T cells. Cells were cotransfected with another GFP-expressing plasmid as a transfection control. **e)** Live-cell imaging of *GFP* mRNA fused to RhoBAST_16_ transcribed under the CAG promoter in HEK293T cells. **f)** Live-cell imaging of the trinucleotide CGG repeat-containing *FMR1-GFP* mRNA fused to RhoBAST_16_ transcribed under the CMV promoter in HeLa cells. **g)** Live-cell imaging of HEK293T cells transfected with a GFP expressing plasmid (negative control) to show the background fluorescence due to the presence of free TMR-DN. At least three independent experiments were carried out with similar results.

**Figure 3.**
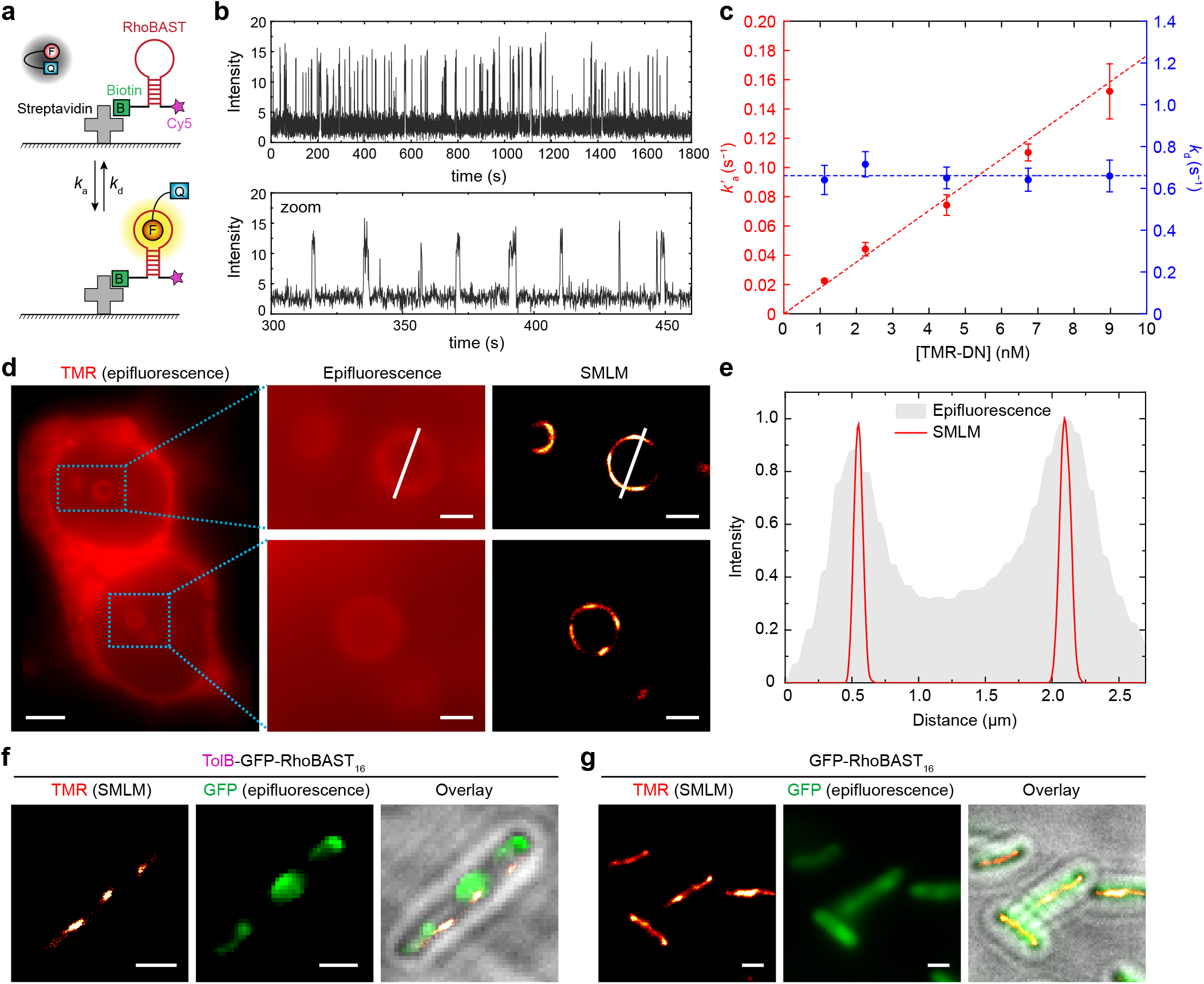
SMLM with the RhoBAST:TMR-DN system. **a)** Scheme showing on and off switching of the fluorescence due to TMR-DN association to and dissociation from RhoBAST immobilized on a cover slip. RhoBAST was modified with biotin at 5′ and Cy5 at 3′ and bound to the glass surface via streptavidin. **b)** Fluorescence time trace taken on an individual, surface-immobilized RhoBAST molecule after addition of TMR-DN (2.2 nM) containing buffer. It reveals intermittent fluorescence emission due to continuous TMR-DN association and dissociation. **c)** Dependence of the average dissociation (*k*_d_ = <*T*_on_>^−1^) and association (*k*’_a_ = *k*_a_ *c* = <*T*_off_>^−1^) rate coefficients on the TMR-DN concentration (*c*). Here, <*T*_on_> and <*T*_off_> are the average times of emission and the lack thereof, respectively. Data points represent mean ± s.d. **d)** Epifluorescence and SMLM images of live HEK293T cells showing hollow spheres containing circular RhoBAST in the nuclei; scale bars, 4 and 1 μm, respectively. **e)** Plot of the fluorescence intensity along the white lines in panel **d**, showing the much higher spatial resolution of SMLM. **f)***tolB-gfp* and **g)***gfp* mRNA fused to RhoBAST_16_ in fixed bacteria was imaged via SMLM and epifluorescence. These images are also shown as overlays on brightfield images. At least three independent experiments were carried out with similar results. Scale bars, 1 μm.

For a FLAP to function in a biological environment, folding into a unique fluorophore-binding structure is essential, as alternative folds would lead to lower sensitivity. To eliminate alternative secondary structures predicted for SRB-3 (Supplementary Fig. 5), we rationally exchanged the U8-A19 base pair by C-G and the UUCG tetraloop by GAAA (Fig. 1e). Indeed, these mutations were beneficial, resulting in a new aptamer dubbed RhoBAST (**Rho**damine **B**inding **A**ptamer for **S**uper-resolution Imaging **T**echniques). It features a fraction of 86 ± 5% in the proper fold, a 26-fold fluorescence turn-on and a *K*_D_ of 15 ± 1 nM (Fig. 1f).

SRB-2, SRB-3 and RhoBAST have similar circular dichroism (CD) spectra (Supplementary Fig. 6). The fluorescence emission of their complexes with TMR-DN is independent of sodium or potassium ions but strictly depends on the magnesium ion (Mg^2+^) concentration (Supplementary Fig. 7), in contrast to G-quadruplex-forming FLAPs^11–13^. Yet, RhoBAST:TMR-DN retains more than 90% of its maximum fluorescence at 0.25 mM Mg^2+^, implying that RhoBAST functions effectively under physiological conditions (Supplementary Fig. 7). RhoBAST:TMR-DN exhibits excellent brightness and well-defined fluorescence spectra, with excitation and emission maxima at 564 and 590 nm, respectively (Fig. 1g,h). It is 4.6- and 7.6-fold brighter than Broccoli:DFHBI-1T^11^ and Corn:DFHO^12^, respectively (Supplementary Table 1). With a melting point, T_m_, of 79 °C, RhoBAST is extremely thermostable. Consequently, upon raising the temperature to 37 °C, RhoBAST:TMR-DN retains 69% of its fluorescence intensity at 25 °C, whereas it drops to 54% for SRB-2:TMR-DN (Supplementary Fig. 6). Furthermore, RhoBAST:TMR-DN appears remarkably photostable; its fluorescence only decreased by about 1% over 3 h under continuous illumination, whereas 12% of the rhodamine-based JF_549_-Halo fluorophore and 50% of the mRuby3 fluorescent protein were photobleached under identical conditions (Fig. 1i). Taken together, these favorable characteristics make the RhoBAST:TMR-DN system a superb genetically encoded tag to image RNA in live cells.

Next, we evaluated how RhoBAST performs in *E. coli* cells in comparison to SRB-2 and SRB-3. Single copies of the aptamers without stabilizing scaffolds were fused to the 3′ untranslated region (UTR) of *gfp* (Fig. 2a). In the TMR channel of confocal images, the signal-to-background (S/B) ratios for bacteria expressing *gfp-RhoBAST, gfp-SRB-3* and *gfp-SRB-2* mRNAs were, respectively, 14 ± 1, 8 ± 1 and 4 ± 1. S/B ratios were calculated as the ratio of the fluorescence intensity (mean ± s.e.m.) at the poles of bacteria expressing *gfp-aptamer* to that of bacteria expressing only *gfp* mRNA (Fig. 2a,b). Moreover, the fluorescence intensity of multiple RhoBAST repeats increased almost linearly up to 16 (Supplementary Fig. 8), which enables imaging of low-abundance RNAs. To analyze the sensitivity of RhoBAST:TMR-DN, bacteria expressing different amounts of *gfp-RhoBAST_16_* mRNA were fixed with paraformaldehyde and imaged. Even at a level of ~1 molecule of *gfp-RhoBAST_16_* per cell (quantified by RT-qPCR), significantly higher fluorescence was detected in the TMR channel than in control bacteria expressing only *gfp* mRNA (Supplementary Fig. 8). Furthermore, to image endogenously expressed mRNAs, multiple RhoBAST repeats were stably introduced into the genome of *E. coli* K12, and *ompA*, *rnaseE* and *cat* mRNAs were visualized displaying distinct localization patterns (Fig. 2c).

To investigate the applicability of RhoBAST:TMR-DN to the imaging of live mammalian cells, a diverse set of RNA species with different reported localization patterns and transcribed from different promoters were examined. First, RhoBAST embedded into the Tornado plasmid system^14^ was expressed as a single-copy, circular aptamer under the control of the U6 promoter in HEK293T cells. Besides nuclear puncta, most of the circular RhoBAST was observed in the cytosol (Fig. 2d), as previously reported for circular Broccoli^14^ and Corn^15^. This demonstrates that RhoBAST folds properly and functions as expected in mammalian cells. Second, *GFP* mRNA tagged with RhoBAST_16_ was expressed under the CAG promoter in live HEK293T cells, revealing the typical cytosolic localization of *GFP* mRNA (Fig. 2e). Third, the CGG trinucleotide repeat-containing *FMR1* (fragile-X mental retardation gene)-*GFP* mRNA fused to RhoBAST_16_ was expressed in HeLa cells under the CMV promoter. It showed the distinct nuclear aggregation pattern reported for this mRNA^16^ (Fig. 2f). The low fluorescence in the TMR channel of control cells demonstrates the specific effect of RhoBAST on the TMR-DN fluorescence (Fig. 2g).

RhoBAST:TMR-DN appears well-suited for SIM^17^ owing to its remarkable photostability and brightness. We examined its performance for SIM by imaging *tolB-gfp-RhoBAST_16_* mRNA in bacteria. The TolB signal peptide causes localization of the mRNA at the inner bacterial membrane due to signal recognition particle (SRP) mediated co-translational translocation^18^ of TolB-GFP. SIM images clearly revealed that *tolB-gfp-RhoBAST_16_* mRNA localized at the inner membrane, whereas *gfp-RhoBAST_16_* lacking the *tolB* sequence or carrying another SRP-independent signal peptide sequence showed substantially different localizations (Supplementary Fig. 9).

Next, we examined the compatibility of RhoBAST:TMR-DN with SMLM by measuring the kinetic parameters of TMR-DN association to and dissociation from the aptamer, generating intermittent fluorescence emission, which is essential for SMLM. To this end, RhoBAST molecules were sparsely immobilized on glass surfaces (Fig. 3a) and, after adding TMR-DN solution, fluorescence intensity time traces from individual molecules were acquired (Fig. 3b). Analysis of the fluorescence on and off times in these traces as a function of the TMR-DN concentration in the range of 1 – 10 nM yields dissociation (*k*_d_ = 0.66 ± 0.01 s^−1^) and association (*k*_a_ = 1.8 ± 0.1 × 10^7^ M^−1^s^−1^) rate coefficients (Fig. 3b,c). These values indicate that a TMR-DN ligand remains bound to RhoBAST for 1.5 s, dissociates and is replaced by a new ligand within about 5 s (at 10 nM). Thus, RhoBAST features a more than two orders of magnitude higher blinking frequency than other FLAPs (Supplementary Table 2). Due to continuous ligand exchange, the number of photons captured from the aptamer is not limited by photobleaching. As a consequence, molecular positions can be determined precisely, as in DNA-PAINT^19^.

To assess RhoBAST:TMR-DN in SMLM applications, we visualized structures inside the cell nucleus formed upon expression of circular RhoBAST in live and fixed HEK293T cells. Intriguingly, we observed hollow spheres in the nucleus, details of which are hardly visible in the epifluorescence image (Fig. 3d, Supplementary Fig. 10). Elaborate immunofluorescence and colocalization studies of these subnuclear structures with literature-reported nuclear bodies (Supplementary Fig. 11) revealed that cyclic RhoBAST in the nucleus co-localized with NONO and PSPC1, proteins that are crucial components of paraspeckle nuclear bodies^20^. Presumably, the high concentration of circular RhoBAST in the nucleus triggers phase separation of RNA with the assistance of NONO and PSPC1, forming paraspeckle-like nuclear bodies (Supplementary Fig. 12).

Multiple repeats of RhoBAST can be introduced to increase the total number of association and dissociation events within a given time interval. Thus, the overall image acquisition time decreases, as photons can be collected at a higher rate to pinpoint the position of the target RNA, albeit with somewhat lower spatial resolution due to the larger tag size. Indeed, rapidly acquired SMLM images of RhoBAST_16_-tagged *gfp, tolB-gfp and phoA-gfp* mRNAs show the correct localization patterns (Supplementary Fig. 13-14), and images reconstructed from 5,000 camera frames (Fig. 3f,g) confirm the excellent image quality.

Here we have introduced a next generation rhodamine-binding aptamer, RhoBAST, as a genetically encoded tag for super-resolution RNA imaging. In comparison to its parent SRB-2, it features markedly improved folding and thermal stability, higher affinity to TMR-DN as well as higher fluorescence quantum yield and photobleaching resistance. Taken together, these advances result in RhoBAST’s excellent performance as an RNA marker for fluorescence imaging. The RhoBAST:TMR-DN system is the first example of a FLAP that is well suited for live-cell SMLM, owing to a continuous and fast fluorophore exchange combined with exceptionally high photostability and brightness. With RhoBAST:TMR-DN, subcellular and subnuclear structures of RNA can readily be visualized with high localization precision in live or fixed specimens.

## Methods

### General

All reagents were purchased from Sigma-Aldrich or Thermo Fisher Scientific unless otherwise specified and used without further purification. Reverse phase HPLC purifications were performed on a Luna® 5 μm C18(2) 100 Å column, 250×10 mm (Phenomenex Ltd.) and compounds were eluted with a mixture of buffer A consisting of 100 mM Et_3_N/AcOH (pH 7.0) in milli-Q-water and buffer B containing 100 mM Et_3_N/AcOH (pH 7.0) in an acetonitrile/water (4:1) mixture. High resolution mass spectra were recorded on a Bruker microTOFQ-II ESI mass spectrometer. DNA and RNA concentrations were determined with a NanoDrop ND-1000 spectrophotometer (NanoDrop Technologies). Fluorescence measurements were performed with a JASCO FP-6500 spectrofluorometer with an ETC-273T digital temperature controller. CD spectra were recorded on a Jasco (model J-810) spectropolarimeter. Agarose gels were stained with ethidium bromide and visualized by UV illumination using an AlphaImager™ 2200 (Alpha Innotech). Absorbance spectra were recorded on a Cary 50 UV-Vis spectrophotometer (Varian). Synthetic DNA oligonucleotides were purchased from Integrated DNA Technologies (IDT). Restriction endonucleases were purchased from Thermo Fisher Scientific. HEK293T (DSMZ, ACC 635) and HeLa cells (DSMZ, ACC 57) were cultured at 37 °C under 5% CO_2_ in Dulbecco’s Modified Eagle’s Medium containing high glucose, HEPES and glutamine without phenol red (Thermo Fisher Scientific) supplemented with 10% FBS, 100 unit/ml penicillin and 100 μg/ml streptomycin. DH5α, K12 and BL21 Star™ (DE3) cells (Thermo Fisher Scientific) were typically grown at 37 °C with shaking at 150 rpm in Luria-Bertani (LB) medium.

### Synthesis

Synthesis and characterization of TMR-SS-Biotin (used for SELEX) and TMR-DN was explained in **Supplementary Note**.

### DNA library preparation for SELEX

A 93-nucleotide long single-stranded DNA oligonucleotide library consisting of 15% doped 54-nucleotide SRB-2 sequence flanked by two constant primer binding sites was synthesized (IDT). DNA library sequence (5′ to 3′): GGAGCTCAGCCTTCACTGC(N3)(N3)(N1)(N1)(N2)(N2)(N4)(N2)(N3)(N2)(N4)(N4)(N2)(N3)(N3)(N2)(N3)(N1)(N4)(N3)(N1)(N4)(N3)(N3)(N1)(N3)(N1)(N3)(N3)(N2)(N3)(N2)(N1)(N1)(N3)(N3)(N4)(N4)(N1)(N1)(N2)(N2)(N3)(N2)(N2)(N4)(N2)(N1)(N3)(N3)(N4)(N4)(N2)(N2)GGCACCACGGTCGGATCCAC. N1, N2, N3 and N4 are mixtures of {85% A, 5% C, 5% G, 5% T}, {5% A, 85% C, 5% G, 5% T}, {5% A, 5% C, 85% G, 5% T} and {5% A, 5% C, 5% G, 85% T}, respectively.

0.5 nmol of the oligonucleotide pool (~3 × 10^14^ molecules) was amplified in a 1 mL PCR reaction for 8 cycles by using the forward (Primer A: 5′-TC<u>TAATACGACTCACTATA</u> GGAGCTCAGCCTTCACTGC-3′) and reverse primers (Primer B: 5’– GTGGATCCGACCGTGGTGCC – 3’) to yield the double-stranded DNA template for the transcription of the library. After phenol:chloroform:isoamyl alcohol (25:24:1, pH 8) extraction, the PCR product was precipitated with sodium acetate and ethanol. The DNA pellet was dissolved in water and directly used for *in vitr*o transcription reaction.

### SELEX

For the first round of selection, a large-scale *in vitro* transcription mixture (1 mL) containing double-stranded DNA template (0.5 μM), transcription buffer (40 mM Tris pH 8.1, 2 mM spermidine, 22 mM MgCl_2_, 0.01% Triton-X-100), DTT (10 mM), BSA (0.01 mg/mL), NTP (4 mM of each ATP, CTP, GTP and UTP) and T7 RNA polymerase (1 μM, lab prepared stock) was prepared. After incubation for 4 h at 37 °C, DNaseI (50 U) was added into the mixture and incubated for 30 min at 37 °C. RNA was purified on a 10% denaturing polyacrylamide gel, excised from the gel and eluted into 0.3 M sodium acetate buffer (pH 5.5) overnight. RNA was precipitated by adding isopropanol and then dissolved in water. To promote the correct folding of the aptamer, the RNA solution was incubated at 75°C for 2 min, and slowly cooled to 25 °C over 10 min. Then, 1/5 volume of 6× aptamer selection buffer (ASB) containing 120 mM HEPES (pH 7.4), 30 mM MgCl_2_ and 750 mM KCl was added and the RNA library was incubated at 25 °C for another 10 min.

TMR-decorated beads were prepared by mixing TMR-SS-Biotin (dissolved in water) with streptavidin-beads (0.2 mL) in 1× ASB. After 5 min, TMR-beads were washed with 1× ASB to remove excess, unbound TMR-SS-Biotin (if any). Next, the folded RNA was mixed with the TMR-beads and shaken for 1 h at 25 °C. To remove unbound and low affinity aptamers, the resin was washed with seven volumes of 1× ASB and the TMR-bound RNA was eluted with 20 mM DTT which cleaves the disulfide bond between TMR and biotin. The eluted RNA was ethanol precipitated with glycogen (20 μg), reverse-transcribed using SuperScript III (Invitrogen), PCR amplified for 10 cycles and the purified double-stranded DNA template was subjected to the next round for selection. Seven rounds of SELEX were performed using the described protocol with progressively increased stringency. The TMR concentration on the streptavidin resin was decreased from 40 μM (round 1) to 10 μM (rounds 2 ‒ 7) and the RNA concentration was decreased from 20 μM (round 1) to 10 μM (rounds 2 ‒ 4) and 5 μM (rounds 5 ‒ 7), whereas the number of washes to remove low affinity binders from the resin was increased to 14 volumes of 1× ASB in the final round. After each round of SELEX, the efficiency of the selection was monitored by quantifying the amount of eluted RNA and measuring the fluorescence enhancements of TMR-DN (25 nM) upon addition of the RNA pool (500 nM). After seven rounds, the double stranded DNA was cloned into a vector, which was subsequently transformed into bacteria, and 96 individual colonies were sent for Sanger sequencing.

### Screening of SRB-2 mutants for fluorescence enhancement

Before fluorescence measurements, SRB-2 mutants (1.2 μM, dissolved in water) were incubated at 75 °C for 2 min, and then cooled to 25 °C within 10 min to promote the correct folding of the aptamer. Then, 1/5 volume of 6× ASB was added and the aptamer solution was incubated at 25 °C for an additional 10 min. The fluorescence intensity of the aptamer solutions after addition of TMR-DN (20 nM) was recorded, and divided by the fluorescence of TMR-DN without any aptamer to yield fluorescence enhancement values of each mutant. The best mutants were further characterized in detail.

### *In vitro* characterization of SRB aptamers

Aptamers were always folded as described above and all properties were measured in 20 mM HEPES (pH7.4), 125 mM KCl and 1 mM MgCl_2_ buffer unless specified. For fluorescence measurements, the excitation and emission maxima of the aptamer:TMR-DN complexes were used. The fluorescence turn-on value is defined as the ratio of the emission peak intensity of the fluorophore (20 nM) in the presence of SRB aptamers (500 nM) to that of the fluorophore (20 nM) in the absence of SRB aptamers.

Dissociation constants (*K*_d_) for aptamer:TMR-DN complexes were determined by measuring the increase in the fluorescence intensity as a function of increasing RNA concentration in the presence of a fixed amount of TMR-DN (1 nM) at 25 °C. Dissociation constants were calculated after fitting the curves to equation 1 by using least-squared fitting (OriginPro 8.5.1):

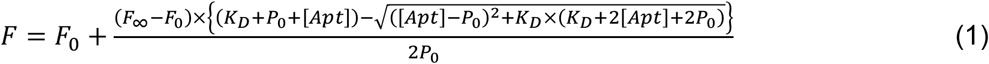

where *F* is the fluorescence at any given aptamer concentration, *F*_0_ is the fluorescence of the free TMR-DN with initial concentration of *P*_0_, *F*_∞_ is the maximum fluorescence intensity reached when all TMR-DN is in complex with the aptamer, [*Apt*] is the final concentration of added aptamer and *K_D_* is the equilibrium dissociation constant.

To determine the Mg^2+^ dependence, 1/5 volume of 6× ASB buffer containing different amounts of MgCl_2_ (0 ‒ 10 mM) was added to the RNA (600 nM in water, already incubated at 75 °C for 2 min and cooled to 25 °C) and incubated for 10 min at 25 °C.

To determine the dependence of the aptamer:TMR-DN fluorescence on monovalent cations (Li^+^, Na^+^, K^+^), 1/5 volume of a 6× buffer containing 750 mM of the corresponding cation as chloride salt or without any monovalent cation, 120 mM HEPES (pH 7.4) and 6 mM MgCl_2_ was added to the aptamer solutions (600 nM in water, already incubated at 75 °C for 2 min and cooled to 25 °C) and incubated for 10 min at 25 °C. Fluorescence intensities were measured upon addition of TMR-DN (500 nM).

To study the temperature dependence of the aptamer:TMR-DN complexes, folded aptamers (500 nM) were incubated with TMR-DN (500 nM) for 10 min. Then, the fluorescence decay was recorded upon increasing the temperature from 25 °C to 65 °C (0.5 °C/min).

To obtain CD spectra, aptamers (5 μM) were folded in 20 mM Na_2_HPO_4_/NaH_2_PO_4_ (pH 7.2), 125 mM KCl and 1 mM MgCl_2_ as previously described and incubated at 25 °C for 10 min. Measurements were performed at 25 °C with a 1 nm resolution using a 1-mm cell. CD melting curves of the aptamers (5 μM) were recorded upon increasing the temperature from 25 °C to 98 °C (2 °C/min) by monitoring the amplitude at 268 nm.

The folding assay was carried out using a slightly modified procedure of Strack *et al* ^21^. Briefly, the fluorescence intensity of a mixture containing 500 nM aptamer (10-fold in excess) and 50 nM TMR-DN was measured (*F*_1_) where the amount of formed complex was controlled by the limiting component TMR-DN. Next, the fluorescence intensity of a mixture containing 50 nM aptamer and 500 nM TMR-DN (10-fold in excess) was measured (*F*_2_), where the amount of formed complex is controlled by the limiting component SRB. Finally, the fluorescence intensity of 500 nM TMR-DN was measured (*F*_3_) for fluorescence background corrections due to unbound excess TMR-DN. Fluorescence intensities were measured in 20 mM HEPES (pH 7.4), 1 mM MgCl_2_ and 125 mM KCl at 25 °C. The fraction of the correctly folded aptamer (*f*) was calculated by using equation 2:

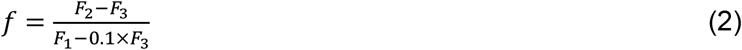

Quantum yields (QYs) of the aptamer:TMR-DN complexes were determined using a cross-calibrated sulforhodamine 101 standard (QY = 1.00 in ethanol). The integral of the fluorescence emission spectra and the absorbance of the samples at the excitation wavelength (528 nm) were measured for at least four different concentrations of the compounds. To avoid inner filter effects in the fluorescence measurements, concentrations with an absorbance below 0.05 were used. The obtained values were plotted in a graph (fluorescence integral vs. absorbance). The slope of the linear fit was compared with that of the reference sample and the QYs were determined using equation 3:

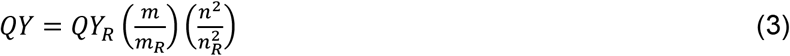

where *m* is the slope, *n* is the refractive index of the solvent and the subscript *R* refers to the reference fluorophore with a known quantum yield.

Photobleaching experiments were performed using a LED setup equipped with a Luxeon Rebel ES Lime LED (nominal wavelength = 567 nm, U = 3.0 V, I = 800 mA). JF_549_-Halo fluorophore^22^ and 50% of the mRuby3^23^ fluorescent protein were prepared in 20 mM HEPES (pH 7.4), 1 mM MgCl_2_, 125 mM KCl, and 0.05% Tween-20 at a final concentration of 20 nM. RhoBAST:TMR-DN was freshly prepared by mixing 500 nM of folded RhoBAST and 20 nM of TMR-DN. All samples were continuously irradiated and the corresponding fluorescence was measured every 20 min over a period of 3 h.

### DNA cloning

For bacterial expression, double-stranded DNA carrying SRB-2, SRB-3 and RhoBAST sequences (**Supplementary Table 3**) flanked by EcoRI and SalI restriction enzyme sites were created in a PCR reaction, double digested and ligated into EcoRI/SalI double digested pET-GFP plasmid (Addgene, Plasmid # 29663) to form pET-GFP-SRB-2, pET-GFP-SRB-3 and pET-GFP-RhoBAST plasmids. The chloramphenycol acetyl transferase (*cat*) gene was PCR amplified from the pLysS plasmid (Thermo Fisher Scientific, isolated from BL21(DE3)pLysS *E. coli*) and cloned into the HindIII and EcoRI sites of pTAC-MAT-Tag2 (Sigma-Aldrich) to yield the pTac-Cat plasmid. A single-stranded, ultramer oligonucleotide containing two repeats of synonymous RhoBAST (**Supplementary Table 3**) flanked by SalI and XhoI sites was synthesized, PCR amplified and blunt-end ligated into PCR-linearized pTac-Cat to yield pTac-Cat-RhoBAST_2_. Another ultramer containing two repeats of synonymous RhoBAST flanked by SalI and XhoI sites was synthesized, PCR amplified, double digested with SalI and XhoI and cloned into XhoI-digested and dephosphorylated pTac-Cat-RhoBAST_2_ to yield pTac-Cat-RhoBAST_4_. A cassette containing four repeats of RhoBAST can be obtained by PCR using pTac-Cat-RhoBAST_4_ as a template. RhoBAST_4_ was then double-digested with SalI and XhoI, and cloned into XhoI-digested and dephosprylated pTac-Cat-RhoBAST_4_ to yield pTac-Cat-RhoBAST_8_. A cassette containing eight repeats of RhoBAST can be obtained by SalI and XhoI double digestion of pTac-Cat-RhoBAST_8_ followed by gel-purification. Cloning this cassette into XhoI-digested and dephosprylated pTac-Cat-RhoBAST_8_ yielded pTac-Cat-RhoBAST_16_.

The GFP gene carrying HindIII at 5′ and SalI-SphI at 3′ was PCR amplified from the pET-GFP plasmid and cloned into HindIII and SphI sites of pTAC-MAT-Tag2 to yield pTac-GFP. A cassette containing 16 repeats of RhoBAST obtained by double digestion and gel purification of pTac-Cat-RhoBAST_16_. It was then cloned into SalI digested and dephosphorylated pTac-GFP to yield pTac-GFP-RhoBAST_16_. TolB (MKQALRVAFGFLILWASVLHA), DsbA (MKKIWLALAGLVLAFSASA) and PhoA (MKQSTIALALLPLLFTPVTKA) signal peptide sequences were fused to the N-terminus of GFP in pTac-GFP-RhoBAST_16_ by PCR. Linearized-plasmids were then phosphorylated and self-ligated to form pTac-olB-GFP-RhoBAST_16_, pTac-DsbA-GFP-RhoBAST_16_, and pTac-PhoA-GFP-RhoBAST_16_.

For mammalian expression, an ultramer ssDNA containing two repeats of synonymous RhoBAST was synthesized, PCR amplified and cloned between SalI and XbaI sites of pAV-U6+27 (Addgene, plasmid #25709) to yield pAV-U6+27-RhoBAST_2_. To generate pAV-Tornado-RhoBAST, an ultramer ssDNA containing RhoBAST sequence embedded into the Tornado system^14^ was synthesized, PCR amplified and cloned between the SalI and XbaI sites of pAV-U6+27. pUC19-CAG-GFP was prepared by digesting the GFP gene from the AAVS1-mEGFP (Addgene, plasmid #91565) plasmid by using SdaI and NotI and cloning it into PCR-linearized pUC19. A cassette containing RhoBAST_16_ was obtained by SalI and XhoI digestion and gel purification of pTac-Cat-RhoBAST_16_. It was then blunted and cloned into pUC19-CAG-GFP which was already digested with the MluI enzyme, blunted and dephosphorylated to yield pUC19-CAG-GFP-RhoBAST_16_. Blunt-end repaired RhoBAST_16_ was cloned into 5′ UTR CGG 99× FMR1-EGFP (Addgene, plasmid #63091) which was digested with the NotI enzyme, blunted and dephosphorylated, to yield 5′ UTR CGG 99× FMR1-EGFP-RhoBAST_16_.

### *E. coli* K12 genome editing

To genomically label *ompA* and *rnaseE* with RhoBAST arrays at the 3′ UTR, double stranded DNA cassettes containing a left-homologous arm, RhoBAST_16_, a terminator, a kanamycin expression system, and a right-homologous arm sequences were prepared by PCR and ligation reactions (**Supplementary Table 4**). On the other hand, the double-stranded DNA cassette for the insertion of *cat* gene into *lacz* locus was prepared by the double digestion of pTac-Cat-RhoBAST_16_ plasmid with BamHI and PagI, gel-purification of the *cat*-RhoBAST_16_ expression system, which was subsequently ligated to a left-homologous arm carrying a synthetic terminator at the 5′-end and to a right homologous arm at the 3′- end (**Supplementary Table 4**). Red/ET homologous recombination in *E. coli* K12 was established using the double stranded DNA casettes prepared as described above and the Quick and Easy *E. coli* Gene Deletion Kit (Gene Bridges) according to manufacturers’ instructions. Selection of bacteria with modified genomes was accomplished in the presence of kanamycin for *ompA* and *rnaseE* or chloramphenicol for *cat*. Successful modifications of the genome were verified by PCR.

### Preparation of 5′-Biotin and 3′-Cy5 labeled RhoBAST

T7 RNA polymerase was added into a transcription mixture containing the double-stranded DNA template (5′- TC<u>TAATACGACTCACTATT</u>**A**GGAACCTCCGCGAAAGCGGTGAAGGAGAGGCGCAAGGTTAACCGCCTCAGGTTCC**AA-3′**, the sequence of T7 φ2.5 promoter is underlined, 1 μM), ATP (0.5 mM), Biotin-HDAAMP (2 mM), CTP (1 mM), GTP (1 mM), UTP (1 mM), DTT (10 mM), spermidine (2 mM), Tris-HCl (40 mM, pH 8.1), MgCl_2_ (22 mM), Triton-X-100 (0.01%), BSA (40 μg/mL) and pyrophosphatase (1 U/mL). After 4 h at 37 °C, the reaction mixture was treated with DNaseI (50 U/mL) for 30 min at 37 °C. RNA was purified by electrophoresis on a 10% denaturing polyacrylamide gel, excised from the gel, ethanol precipitated and dissolved in water. The percentage of biotinylation (~30%) was determined by streptavidin-shift assay. 3′ of Biotin-RhoBAST was further ligated to Cy5, by incubating the RNA (10 μM) with ATP (1 mM), pCp-Cy5 (20 μM) and T4 RNA ligase at 4 °C overnight. RNA was then extracted with phenol:chloroform:isoamyl alcohol (25:24:1, pH 4.5), precipitated using sodium acetate and ethanol, dissolved in water and stored at −20 °C. The percentage of Cy5 labeling (~60%) was calculated by measuring the absorbtion of the Cy5 labeled RNA at 648 nm (extinction coefficient of Cy5 at 648 nm is 250,000 M^−1^cm^−1^).

### Live-cell confocal imaging of bacterial cells

BL21 Star™ (DE3) competent *E. coli* cells were transformed with either pET-GFP (control plasmid) or pET-GFP-SRB-2, -3 or pET-GFP-RhoBAST plasmids. Next day, single colonies were picked from LB-agar/Kanamycin (30 μg/mL) plates and grown in 5 mL of LB medium containing kanamycin (30 μg/mL) overnight at 37 °C with shaking at 150 rpm. A fresh culture with an OD_600_ of 0.01 was started using the overnight culture as a starter culture in a 50 mL falcon flask containing 10 ml of LB medium with 30 μg/μL kanamycin. When OD_600_ was 0.4, IPTG (at desired concentration, ≤1 mM) was added into the culture, and flasks were shaken for an additional 2 ‒ 3 h at 37 °C. Then, 200 μL of the culture was removed, spun down and resuspended in 1 mL of M9 medium. 200 μL of this suspension was transferred into a poly-D-lysine coated 8-well glass chamber and incubated at 37 °C for 20 min. Finally, the wells were gently washed twice and bacteria were incubated with 100 nM of TMR-DN in M9 medium at 37 °C. Bacteria were imaged after 20 min of incubation at 37 °C.

Images were taken on a point-scanning confocal microscope with hybrid scanner (galvo/resonant) equipped with a Nikon N Apo 60× NA 1.4 λs OI (WD 0.14 mm, FOV 0.21 × 0.21 mm^2^) objective. A 405 nm laser was used to excite Hoechst 33342 (detection via an emission filter set of 450 ± 25 nm), a 488 nm laser was used to excite GFP (emission filter set 525 ± 25 nm) and a 561 nm laser was used to excite TMR (emission filter set 595 ± 25 nm). The laser settings were optimized for each condition. Z-stack images were taken with a step size of 250 nm. Images were analyzed by Fiji/ImageJ and the background correction was done by subtracting the mean fluorescence intensity of a surface area without attached *E. coli* cells from the whole image. The mean fluorescence intensity was determined from defined areas at the poles of the bacteria.

For imaging GFP mRNA using pTac-GFP (control plasmid), pTac-GFP-RhoBAST_16_, pTac-TolB-GFP-RhoBAST_16_, pTac-DsbA-GFP-RhoBAST_16_ and pTac-PhoA-GFP-RhoBAST_16_ plasmids, DH5α cells were used and the LB growth medium was supplemented with ampicillin (100 μg/mL) instead of kanamycin.

For imaging *ompA* and *rnaseE* mRNA in *E. coli* K12, LB growth medium was supplemented with kanamycin (15 μg/mL). For imaging *cat* mRNA in *E. coli* K12, LB growth medium was supplemented with chloramphenicol (10 μg/mL).

### Confocal imaging of fixed bacterial cells

700 μL bacteria expressing RNA of interest fused to RhoBAST in LB or M9 medium (as described above) were quickly mixed with 100 μL 8% paraformaldehyde (PFA) solution, and incubated for 10 min at room temperature. Bacteria were then pelleted by centrifugation, washed twice with 1 mL MgPBS (DPBS containing 1 mM MgCl_2_), resuspended in 1 mL MgPBS and stored overnight at 4 °C. On the next day, 200 μL of the culture was transferred into a poly-D-lysine coated 8-well glass chamber and incubated at room temperature for 20 min. Finally, the wells were gently washed twice and bacteria were incubated with 1 μg/mL Hoechst 33342 and 50 nM TMR-DN in MgPBS (or M9 medium) at 37 °C for 20 min. Bacteria were then imaged at room temperature.

### Quantitative PCR with reverse transcription (RT–qPCR)

Chemically competent DH5α competent *E. coli* cells (Thermo Fisher Scientific) were transformed with pTac-GFP (negative control) and pTac-GFP-RhoBAST_16_. On the next day, single colonies were picked from the LB-agar/ampicillin (100 μg/mL) plates and grown overnight in 5 mL of LB medium containing ampicillin (100 μg/mL) at 37 °C with shaking. A fresh culture with an OD_600_ of 0.01 was started using the overnight culture as a starter in a 50 mL conical falcon flask containing 15 mL of LB medium with 100 μg/μL ampicillin. After 3 h (OD_600_ ~0.4), the culture was equally divided into 3 different falcon flasks and different amounts of IPTG (0, 10 μM, 40 μM) were added into each culture flask. Cells were shaken for another 2 h at 37 °C. Then, 2 mL of the cultures were spun down, LB medium was carefully removed from the bacteria pellet and the pellet was resuspended in 450 μL of M9 medium. 400 μL of this mixture were used for total RNA isolation and 50 μL were used for fixed-cell imaging. Total RNA from the bacteria was isolated using RNAzol^®^RT according to the manufacturers’ instructions and obtained RNA pellet was finally dissolved in 200 μL of water.

Total RNA (20 μL, ~2.6 μg) was then subjected to DNaseI (50 U/mL) treatment in 60 μL total volume at 37 °C for 1 h and the DNaseI was inactivated by adding 4 μL of 50 mM EDTA and incubating at 65 °C for 10 min. cDNA was produced using 5 μL of each RNA sample and SuperScript™ IV-RT (Invitrogen) according to manufacturers’ instructions. Reverse-transcription reactions were diluted 10-fold with water prior to qPCR which was performed in a Light Cycler 480 instrument (Roche) using the Brilliant III Ultra-Fast SYBR® Green qPCR Mastermix (Agilent). Forward and reverse GFP primers are 5′-TGCTGCTGCCCGACAACCAC-3′ and 5′-CGGTCACGAACTCCAGCAGGAC-3′, respectively. qPCR data were analyzed with the LightCycler 480 software. As negative controls, no-RT and no-template reactions were used. All reactions were performed in triplicate on the 20 μL scale. 16S rRNA was used as an internal reference gene. For absolute quantification of RNA copy numbers, a small fragment (~180 nucleotide) of GFP gene was *in vitro* transcribed and used as a reference RNA in RT-qPCR.

### Confocal imaging of live mammalian cells

HeLa or HEK293T cells were cultured at 37 °C under 5% CO_2_ in Dulbecco’s Modified Eagle’s Medium (with 4.5 g/L high glucose, 4 mM L-glutamine, 25 mM HEPES and without phenol red) supplemented with 10% FBS, 100 unit/mL penicillin and 100 μg/mL streptomycin. To improve the adherence of HEK293T cells, glass slides were coated with poly-D-lysine before plating the cells.

For imaging experiments, 3 × 10^4^ cells were seeded overnight in μ-Slide chambered coverslips with 8-wells containing 300 μL of medium each. The following day, cells were transfected with the appropriate plasmid using the FuGeneHD transfection reagent (Promega) according to the manufacturer’s protocol. After 48 h at 37 °C, the medium was exchanged with Leibowitz (L15) medium containing 1 μg/mL Hoechst 33342 and 100 nM of TMR-DN. After incubation for 1 h, cells were imaged at 37 °C in a live-cell imaging chamber with controlled humidity.

Images were taken as described for bacterial cells above, except for the following modifications. Z-stack images were taken with a step size of 500 nm. Images were analyzed by Fiji/ImageJ and the background correction was done by subtracting the mean fluorescence intensity of a surface area without adherent cells from the whole image.

### SIM imaging of bacterial cells

DH5α cells *E. coli* cells were transformed with either pTac-GFP (control plasmid) or pTac-GFP-RhoBAST_16_, pTac-TolB-GFP-RhoBAST_16_ or pTac-PhoA-GFP-RhoBAST_16_ plasmids. On the following day, single colonies were picked from LB-agar/Ampicillin (100 μg/mL) plates and grown in 5 mL of LB medium containing ampicillin (100 μg/mL) overnight at 37 °C with shaking at 150 rpm. A fresh culture with an OD_600_ of 0.01 was started using the overnight culture as a starter culture in a 50 mL falcon flask containing 10 mL of LB medium with 100 μg/μL ampicillin. When OD_600_ was 0.4, IPTG (40 μM) was added into the culture, and flasks were shaken for an additional 2 ‒ 3 h at 37 °C. Then, 200 μL of the culture was removed, spun down and resuspended in 1 mL of M9 medium. At this point cells can be fixed using PFA (as described before) and imaged later. For live-cell imaging, 200 μL of this suspension was transferred into a poly-D-lysine coated 8-well glass chamber and incubated at 37 °C for 20 min. Finally, the wells were gently washed twice and bacteria were incubated with 50 nM of TMR-DN in M9 medium at 37 °C. Bacteria were imaged after 20 min of incubation at 37 °C.

Images were taken on a Nikon N-SIM system equipped with total internal reflection fluorescence Apochromat 100 × 1.49 NA oil immersion objective and an electron-multiplying charge-coupled device camera with single-molecule sensitivity (iXon3 DU897E; Andor Technology). A 488 nm laser was used to excite GFP (detection via an emission filter set of 520/45 nm), a 561 nm laser was used to excite TMR (emission filter set 610/60 nm). The laser settings were optimized for each condition. Images were taken sequentially within a small z-stack (step size of 150 nm). Subsequently, reconstruction of the super-resolution images was performed with the NIS imaging and image analysis software (Nikon).

### Imaging of immunostained mammalian cells

3 × 10^4^ HEK293T cells were seeded overnight in poly-D-lysine-coated μ-Slide (chambered coverslip) with 8 wells containing 300 μL of medium. On the following day, cells were transfected with pAV-Tornado-RhoBAST plasmid (~200 ng/well) using FuGeneHD transfection reagent (Promega) according to the manufacturer’s protocol. After 48 h at 37 °C, the medium was exchanged with 300 μL L15 medium. Then, half of the medium was removed, 150 μL 8% PFA (in DPBS, pH 7.4) was added and the solution was gently but quickly mixed. After 10 min at room temperature, cells were washed 3 times with 500 μL of MgPBS. In order to permeabilize the cells, 400 μL of 0.1% Triton X-100 in MgPBS was added into the wells and cells were incubated for 15 min at room temperature. Triton-X-100 was removed and the cells were washed 3 times with 500 μL MgPBS. Before proceeding to immunostaining, cells were incubated with 500 μL of 2% BSA in MgPBS overnight at 4 °C for blocking. On the next day, the appropriate concentration of primary antibody diluted in 500 μL of 0.1% BSA was added to the cells and incubated for 1 ‒ 2 h at room temperature. All antibodies used in this study and their dilution factors are listed in **Supplementary Table 5**. The primary antibody solution was removed from the wells and the cells were washed three times with 500 μL of MgPBS for 5 min. Then, the appropriate concentration of the Alexa488 fluorophore-labeled secondary antibody diluted in 500 μL of 0.1% BSA was added to the cells and incubated for 1 h at room temperature. Finally, the secondary antibody solution was removed from the wells and the cells were washed three times with 500 μL of MgPBS for 5 min. Before imaging, the solution was exchanged with MgPBS containing 1 μg/mL Hoechst 33342 and 100 nM of TMR-DN and the cells were incubated for 10 min at room temperature.

Images were taken as described before for live mammalian cells; imaging conditions identical to those of GFP were used for Alexa488.

### Single-molecule studies of RhoBAST binding kinetics

A home-built total internal reflection fluorescence (TIRF) microscope based on an inverted microscope (Axiovert 200, Zeiss, Göttingen, Germany)^24^ was used. Light from a 638 nm laser (MLD 638, Cobolt AB, Solna, Schweden) and a 561 nm laser (Jive, Cobolt AB) is combined via dichroic mirrors into an acousto-optical tunable filter (AOTFnC-400.650, A-A Opto-Electronic, Orsay, France) for precise and fast control of the laser intensities. The light is guided into a Pellin-Broca prism located on top of a quartz glass sample slide and totally reflected at the boundary between the sample slide and the aqueous sample solution. To ensure undisturbed guidance of the beams to the boundary layer, immersion oil (*n* = 1.518, Immersol 518F, Zeiss) is used between the prism and the quartz glass sample slide. Fluorescence emission is collected through a water immersion objective (C-Apochromat 63×/1.2 W Corr M27, Zeiss). Excitation light in the detection path is blocked by 561 and 635 nm notch filters (Thorlabs, Newton, NJ) and the fluorescence emission is detected by an EMCCD camera (Ixon EM+ DU-897, Andor, Belfast, UK) with a pixel size of 120 × 120 nm^2^.

In order to prepare the sample slides, 75 × 75 × 1 mm^3^ quartz slides (Finkenbeiner, Scientific Glass Blower, Waltham, MA) were sonicated for 30 min at 60 °C in 1 M KOH. This step was repeated with fresh 1 M KOH solution. Afterwards, they were sonicated for 15 min at 60 °C in Millipore water. This step was repeated after water exchange. Then, the slides were dried under nitrogen and plasma-cleaned for 30 min, and incubated overnight in 0.05% (v/v) dichlorodimethylsilane (Sigma-Aldrich, St. Louis, MO) solution in n-hexane (VWR, Radnor, PA). On the following day, the slides were sonicated in *n*-hexane at room temperature for 1 min three times in succession, each time with fresh *n*-hexane.

Cover slips (20 × 20 × 0.17 mm^3^, Menzel Gläser, Thermo Fischer, Waltham, MA) were prepared in essentially the same way, except that plasma cleaning was omitted. Quartz glass slides and cover slips were stored at 4 °C until use.

The quartz slides and cover slips were glued together using two pieces of double-sided adhesive tape, leaving a 3 mm wide channel in the middle. The channel was filled with 20 μL of a 0.01 mg/mL neutravidin solution in PBS and incubated for 10 min. Then, the channel was washed with 100 μL PBS and incubated for 10 min with 100 μL of a 0.1 mg/mL solution of Lutensol AT50 (BASF SE, Ludwigshafen, Germany) in water for surface passivation. The channel was washed again with 100 μL of PBS. To ensure a proper folding of the aptamer, the RhoBAST stock solution (20 nM in water) was heated to 75 °C for 2 min and then slowly cooled to room temperature prior to filling it into the channel. To obtain a sparse decoration of the surface with aptamers, the Cy5-labelled, biotinylated RhoBAST stock was diluted to a final concentration of 25 pM with 1×ASB. The channel was flushed with 100 μL of this solution and incubated for 10 min. Then, the channel was washed with 100 μL ASB and filled with 20 μL of imaging buffer containing the TMR-DN dye at the desired concentration, an enzymatic oxygen scavenging system (1.4 mg/mL glucose oxidase, 0.03 mg/mL catalase, 10% (w/w) glucose), 1 mM Trolox as triplet state quencher and 0.1% (v/v) Tween 20 in 1×ASB. The channel was sealed with transparent nail polish after adding the imaging solution and measurements were started after 10 min to ensure equilibration.

To locate RhoBAST molecules, 100 camera frames (100 ms dwell time) were recorded with 638 nm excitation (7 mW in front of the prism) to excite the Cy5 marker attached to the RhoBAST molecules. Then, the Cy5 was photobleached using 49 mW laser power to avoid energy transfer from bound TMR-DN dyes to the Cy5 label in the subsequent measurement. Afterwards, the TMR-DN dyes were imaged with 561 nm excitation, taking 18,000 frames (100 ms dwell time) with a laser power between 2 and 7 mW.

Images and time traces were analyzed in a semi-automatic way using our own software written in MATLAB 2019a (MathWorks, Natick, MA). Aptamer locations were identified as local intensity maxima in the images taken in the Cy5 channel. Only maxima with an intensity of 80% or more of the global maximum were included. The intensities in the TMR-DN channel were integrated within a 5 × 5 pixels window around the aptamer positions for all frames and combined in intensity time traces. From these traces, on-times (*τ_on_*) and off-times (*τ_off_*) were analyzed by setting a threshold of 3× the standard deviation above the mean background. For each time trace, on- and off-times were fitted with an exponential decay to obtain average *τ_on_* and *τ_off_* values at a single aptamer position. An error of ± 50 ms was assigned to these values to account for the camera dwell time. Average <*τ_on_*> and <*τ_off_*> values taken over different aptamer positions were obtained (for identical TMR-DN concentration) by taking a weighted average according to the uncertainties of the average *τ_on_* and *τ_off_* values of the individual aptamers. Average rate coefficients of dissociation (*k*_d_ = <*T*_on_>^−1^) and association (*k*’a = *k*_a_ *c* = <*T*_off_>^−1^) were calculated. The final *k*_d_ was obtained by averaging over all concentrations and *k*_a_ by a linear regression to the concentration dependence (Fig. 3c). Data were taken at 1.1, 2.2, 4.5, 6.7 and 9.0 nM TMR-DN on 12 (410), 11 (610), 13 (1230), 8 (1000) and 10 (1730) individual molecules, respectively. The numbers in parentheses denote the total number of on-switching events.

### SMLM imaging

A home-built widefield microscope based on an Axio Observer Z1 frame (Zeiss, Göttingen, Germany) with single-molecule sensitivity was used for SMLM, with small modifications from the design described earlier^25^. Light from a 473 nm laser (Gem 473, Laser Quantum, Konstanz, Germany) and a 561 nm laser (Gem 561, Laser Quantum) is combined via dichroic mirrors and passed through an AOTF (AOTFnC-400.650, A-A Opto-Electronic) to ensure precise and fast control of the laser intensities. Two achromatic lenses with focal lengths of 10 and 100 mm (Thorlabs) expand the beam after the AOTF. For widefield illumination, the beam is focused onto the back focal plane of a high numerical aperture oil immersion objective (α Plan-Apochromat 63×/1.46 Oil Corr M27 TIRF, Zeiss) by a scan lens. Fluorescence emission is collected through the same objective and filtered by a quad-band dichroic mirror (z 405/473/561/640, AHF, Tübingen, Germany). After passing a tube lens, a beam splitter (OptoSplit II, Cairn Research, Kent, UK) separates the fluorescence into two images of different color, which are imaged side by side on an EMCCD camera (Ixon Ultra X-7759, Andor, Belfast, UK) with a pixel size of 109 × 109 nm^2^.

Fixed DH5α bacteria expressing *gfp* (control), *gfp-RhoBAST_16_*, *tolB-gfp-RhoBAST_16_*, *dsbA-gfp-RhoBAST_16_* and *phoA-gfp-RhoBAST_16_* were prepared and immobilized on a poly-D-lysine coated 8-well glass chamber as described before. Bacteria were incubated with 15 nM TMR-DN in M9 medium for 30 min and imaged at room temperature. For the measurements, GFP was excited with the 473 nm laser; its emission was collected through a 525/45 nm (center/width) filter. 300 camera frames were collected using continuous illumination with an exposure time of 100 ms per frame and a laser power of 50 μW. Then, the excitation was swiftly changed to 561 nm by means of the AOTF, and TMR-DN was imaged using a 607/70 nm emission filter. 5,000 ‒ 10,000 frames were collected with an exposure time of 100 ms each. The laser power was adjusted in the range 4.9 ‒ 20.4 mW to achieve temporally well-separated blinking events in different cell regions.

To reduce noise in the GFP channel, 300 camera frames were averaged by using Fiji/ImageJ software. For registration of the SMLM data in the TMR-DN channel, custom-written a-livePALM software was used^10^. The major steps of the algorithm include background estimation and Gaussian noise filtering. Based on the detected background information, regions with local maxima are identified; their location is precisely determined with the maximum likelihood estimator (MLE) algorithm^26^. Localization accuracies of the SMLM images are calculated as described in Ref ^27^.

Sample drift during data acquisition was corrected by a cross-correlation based analysis ^28^. A sub-stack of 500 frames was selected to reconstruct a reference image. The following sub-stack was reconstructed and its drift with respect to the reference frame was determined by cross-correlation and compensated. The combined image was subsequently taken as the new reference image. This procedure was repeated for all sub-stacks of the total image stack.

### SMLM imaging of mammalian cells

HEK293T cells expressing Tornado-RhoBAST were prepared as described before. For imaging live cells, 48 h after the transfection, the medium was exchanged by L15 medium containing 100 nM TMR-DN. After incubation for 1 h, cells were imaged at 37 °C and 5% CO_2_ in a live-cell imaging chamber. TMR-DN was imaged with 561 nm laser excitation (3.1 mW), the fluorescence was detected through a 607/70 nm emission filter. 1000 frames were collected with an exposure time of 30 ms.

For imaging fixed cells, the cells were incubated in DPBS medium containing either 10 nM or 20 nM TMR-DN and 1 mM MgCl_2_. After incubation for 1 h, cells were imaged at room temperature with 561 nm excitation and a 607/70 nm emission filter. 5000 frames were collected with camera dwell times between 15 and 50 ms and laser powers between 3.1 mW and 14.6 mW to achieve well separated blinking events in each case.

The data were analyzed with the a-livePALM software^10^ as described above; no drift correction was applied. The final images were prepared with Fiji/imageJ; intensity profiles were analysed with Fiji/imageJ using a line width of five pixels at corresponding positions in the epifluorescence and SMLM images.

## Acknowledgements

M.S. and A.J. were supported by the Deutsche Forschungsgemeinschaft (DFG Grant # Ja794/11-1) and G.U.N. by the Helmholtz Association (Program Science and Technology of Nanosystems) and by the DFG (GRK 2039). We thank the Nikon Imaging Center, Heidelberg for granting access to their facilities and Dr. Ulrike Engel for technical advice in fluorescence microscopy.

## Author contributions

M.S., G.U.N. and A.J. designed the study. A.M. and M.S. evolved RhoBAST. D.E. and M.S. characterized RhoBAST. M.S. created all plasmid constructs and strains. M.S carried out confocal and SIM microscopy. J.L. and B.H. established single-molecule binding kinetics experiments. J.L. carried out SMLM. M.S., K.N., G.U.N. and A.J. supervised the work. M.S. wrote the first draft and all authors contributed to reviewing, editing and providing additional text for the manuscript.

## Additional information

**Supplementary information** is available for this paper.

