## Supplementary Information (SI) for "RhoBAST - a rhodamine-binding aptamer for super-resolution RNA imaging"

| Clone | Sequence | Length (nt) | Freq | Turn-on |
|---|---|---|---|---|
| SRB-2 | ------------GGAACCTCG-CTTCGGCGATGATGGAGAGGCGCAAGGTTAACCGCCTC-AGGTTCC--------------- | 54 | - | 17 |
| 2 | -----------GATGCCT--GGCTTCGGCGAGGAGGGAGGGACGGGAGGATATTCGCCCC-AAGTTAC--------------- | 54 | 1 | 4.0 |
| 86 | -----------GGTGCCT--GGCTTCGGCGAGGAGGGAGGGACGGGAGGATATTCGCCCC-AAGTTAC--------------- | 54 | 1 | 1.7 |
| 89 | -----------GGTACCT--GGCTTCGGCGAGGAGGGAGGGACGGGAGGATATTCGCCCC-AAGTTACGCGGCCGCGGGTCG- | 68 | 1 | 1.9 |
| 14 | -----------GGTGCCTTCGTT-TCGGCAAGGAGGAAGAGGCCCCAGGTTAATCGCCTC-GGGTTTC--------------- | 55 | 7 | 1.6 |
| 40 | ------------GTCCCTTCGGCGTTGGCAAGGA------AACGCAAGGTTAATAGCCTC-GGGATGT--------------- | 49 | 1 | 1.3 |
| 85 | ------------GTCCCTTCGGCGTTGGCAAGGA------AACGCAAGGTTAATAGCCTC-GGGATGC--------------- | 49 | 1 | 1.5 |
| 52 | ------------GTCCCTTCGGCGTTGGCAAGGA------AACGCGAGGTTAATAGCCTC-AGGATGC--------------- | 49 | 1 | 3.4 |
| 3 | -----------GTAAGCTTCA-CTTCGGCAAGGAGGAG-AGTGGCAAGGTTAAGCACCTC-AGGATCA--------------- | 54 | 2 | 1.5 |
| 6 | ------------AGACTTTCG-CTTCGGCAAGGAGGAGGAGGCGCAAGATTAAGCACCTC-AGTTTGC--------------- | 54 | 5 | 1.5 |
| 30 | ------------GGAGTTTCG-CTTCGGCAAGGAGGAGCAG-CGCAATGCTGTACGCCGC-AGTTTTC--------------- | 53 | 2 | 1.6 |
| 9 | ------------GGAGTTTCG-CTTCGGCAAGGAGGAGCAG-CGCAATGTTGTACGCCGC-AGTTTTC--------------- | 53 | 1 | 1.4 |
| 11 | ------------GTAACTTCG-CTTCGGCAAGGAGGAGATG-AGGAACGTCAACGGCCTC-ATGTTAC--------------- | 53 | 4 | 1.6 |
| 19 | ------------GGAGTTTCG-CTTCGGCAAGGAGGAGATG-AGGAACGTCAACGGCCTC-ATGTTAC--------------- | 53 | 1 | 3.7 |
| 44 | ------------CGCACCTCG-TTTCGGCGACGGCAAGGAGTCGCGAGGTTAACCGCCTC-ACACTGC--------------- | 54 | 3 | 1.5 |
| 80 | ------------GGAACGTTG-GTTCGGCGACGGCAAGGAGTCGCAAGGTTCACTTCCTC-ATGTTCC--------------- | 54 | 1 | 4.6 |
| 92 | ------------GTAACGCAG-CTTCGGCTCTGGCAAGGAAGAGCAAGGTTGACTGCCTC-ACGGTCC--------------- | 54 | 1 | 1.1 |
| 37 | ------------TGAACCTCG-CTTCTGCCCTGATCGAGAGGGGCAATGCAAACC-TCTC-AGGTTCC--------------- | 53 | 5 | 1.5 |
| 4 | ------------TGAACCTCG-CTTCTGCCCTGATCGAGAGGGGCAATGCAAACC-TCTC-AAGTTCC--------------- | 53 | 1 | 1.5 |
| 96 | ------------TGAACCTGG-CTTCCGCGTTGATGCCGCGGCGCAAGTTTAACCGCCAC-AGGTTAC--------------- | 54 | 1 | 1.5 |
| 59 | ------------GGAACCTCG-CTTCGACGACAATGGTATCGAGCAAGATTATCTGACTC-AAGTTGA--------------- | 54 | 1 | 1.2 |
| 69 | ------------GCAACGTCG-CTTCAGTGATGA-----------AGGA-----------------GA--------------- | 27 | 1 | 1.1 |
| 64 | ------------GGAGCGTCG-CCGCAGTGGTGATGGAGAGGCCCAAGGTTAACGGCCTC-AGTGTAA--------------- | 54 | 1 | 5.2 |
| 1 | ------------TAATCCTCG-GTTCGGTGATGAAGGAGCGGCACAAGGTTAACTGCCGC-AGGCTTG--------------- | 54 | 1 | 11.2 |
| 95 | ------------AGAACCGCT-CTGCGGTGATGAAGGAGCGGTGCAAGGTTAACCACCGC-AGGCTCG--------------- | 54 | 1 | 9.6 |
| 15 | ------------TAAATCGCG-ATTCGGTGATGATGGACATGCGCAAGGTTAACCGCATG-AGGTTAG--------------- | 54 | 3 | 6.6 |
| 5 | ------------AAAAGCTCG-CTGCGGTGATGATGGACATGCGCAAGGTTAACCGCATG-AGGTTAG--------------- | 54 | 1 | 6.7 |
| 34 | ------------TAAGACTCG-TTTCGGCGATGATGGAGAGGCGCAAGGTTAACCGCCTCGAGGTCTG--------------- | 55 | 1 | 3.4 |
| 50 | ------------TAAGACTCG-CTACAGCGATGATGGAGGGGNGCAACGTTAACCGCCTCGAGGTCTG--------------- | 55 | 1 | 2.6 |
| 74 | ------------GCACCCACG-CTGCGGCGATGATGGAGAAGAGCAAGGTTAACCTCTTC-AGGTTGT--------------- | 54 | 1 | 8.8 |
| 75 | ------------GTAAACACG-CTGCGGCGGTGATGGAGAGGCACAAGGTTAACTGTCTC-AGGTTGT--------------- | 54 | 1 | 7.7 |
| 22 | -----------GGCAAGGA-G-CATGGGAGATGATGGAGAGACGCAAGTTCAACATCCTC-CGGTTTG--------------- | 54 | 1 | 1.6 |
| 18 | ------------GGAACCTCG-ACTCGGCGGAGATAGAGCGGCGGGAGGGTTACCGGGTC-ATGTTCT--------------- | 54 | 2 | 1.4 |
| 70 | ------------TGAACCTCG-ACCCGGTGATGATGAACGGACGCGACGTTGACCGCCTC-GGTTTTC--------------- | 54 | 1 | 1.6 |
| 16 | TCTCCGTGCAATCCTACCTTAACTTCTGCTTCGGCAGAGAGGAGCCA--TTGGTAGCTCC-GC--TTT--------------- | 63 | 4 | 1.5 |
| 91 | TCTCCGTGCAATCCTACCTTAACTTCTGCTTTGGCAGAGAGGAGCCA--TTGGTAGCTCC-GC--TTT--------------- | 63 | 2 | 1.3 |
| 28 | ---------AATCCGACCTTG---TCTGCTTCGGCAGAGAGGAGCCA--CTGGTAGCTCC-GC--TTT--------------- | 51 | 1 | 4.8 |
| 94 | --------GAAACCTACCT-----ACGGCGATGGTTGAGGTGCCCTAGGTTAACCGTCTC-GCGTTTT--------------- | 54 | 1 | 1.5 |
| SRB-2 | ------------GGAACCTCG-CTTCGGCGATGATGGAGAGGCGCAAGGTTAACCGCCTC-AGGTTCC--------------- | 54 | - | 17 |

**Supplementary Figure 1. Screening SRB-2 mutants for fluorescence turn-on.** Sanger sequencing results of the selected SRB-2 mutants, their frequencies and fluorescence turn-on values. Sequences recovered from Sanger sequencing were PCR amplified using primers A and B binding to the constant regions of the library. Then, they were transcribed into RNA using T7 RNA polymerase, extracted and precipitated. Without further purification, all mutants (~500 nM) were incubated with TMR-DN (25 nM) for determination of fluorescence turn-on values. Mutants with a turn-on factor >6 were further evaluated.

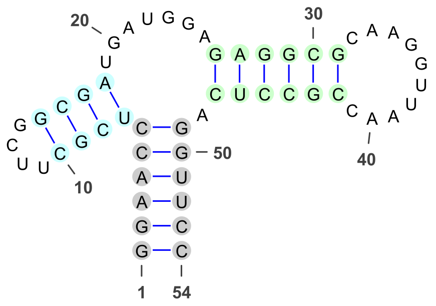


SRB-2

| Clone | Truncated sequence | Turn-on | Change |
|---|---|---|---|
|  | ...... .... .... ...... ...... ...... |  |  |
| SRB-2 | GGAACC UCGC UUCG GCGA UGAUGGA GAGGCG CAAGGUUAAC CGCCUC A GGUUCC | 17 | - |
| 1-v2 | GGAACC UCGG UUCG GUGA UGAAGGA GCGGCA CAAGGUUAAC UGCCGC A GGUUCC | 9.3 | -17% |
| 95-v2 | GGAACC GCUC UGCG GUGA UGAAGGA GCGGUG CAAGGUUAAC CACCGC A GGUUCC | 2.8 | -71% |
| 74-v2 | GGAACC ACGC UGCG GCGA UGAUGGA GAAGAG CAAGGUUAAC CUCUUC A GGUUCC | 8.1 | -8% |
| 75-v2 | GGAACC ACGC UGCG GCGG UGAUGGA GAGGCA CAAGGUUAAC UGUCUC A GGUUCC | 7.8 | 0 |
| 5-v2 | GGAACC UCGC UGCG GUGA UGAUGGA CAUGCG CAAGGUUAAC CGCAUG A GGUUCC | 1.5 | -78% |
| 15-v2 | GGAACC GCGA UUCG GUGA UGAUGGA CAUGCG CAAGGUUAAC CGCAUG A GGUUCC | 3.4 | -48% |

**Supplementary Figure 2. Effect of sequence truncation on the fluorescence turn-on values of the best mutants.** The primer binding sites at the 5′ and 3′ ends of the aptamers were truncated and the terminal stem of the mutants was replaced with the one found to promote the correct folding of SRB-2. Truncated clones, represented as “Clone#-v2”, were transcribed and their fluorescence turn-on values were determined. Comparison of these values with the original ones indicates that primer-binding sequences play a significant role in the overall folding.

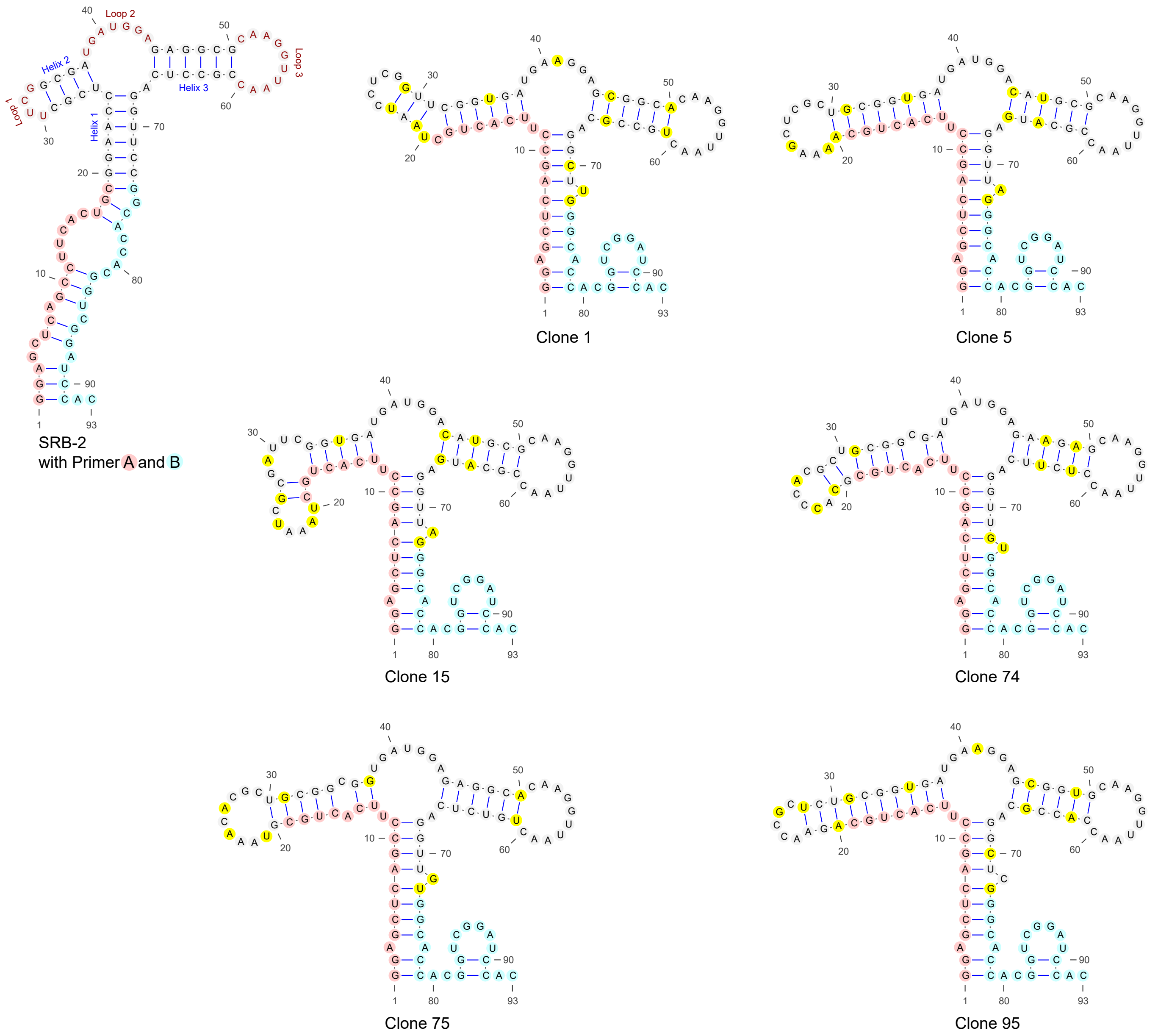


**Supplementary Figure 3. Primer A is crucially involved in the folding of mutants with the highest fluorescence turn-on.** Secondary structure predictions and fluorescence turn-on values show that Primer A has an important role in the correct folding of the selected mutants. Primer A is also responsible for the insertion of a single U nucleotide between C6 and U7 in the SRB-2 sequence. Collectively, these analyses revealed that *i*) the UUCG tetraloop (loop 1) is not required for activity and the length of helix 2 (H2) is variable, *ii*) the sequences of helices H1, H2 and H3 are quite flexible, *iii*) the sequences of loops 2 and 3 is strictly conserved among the active mutants. The mutants with the highest fluorescence turn-on, namely clone 1 and clone 95, had a single U nucleotide insertion between C6 and U7 as well as a U22A mutation in SRB-2. These mutations were taken into account in the design of the new aptamer, SRB-3.

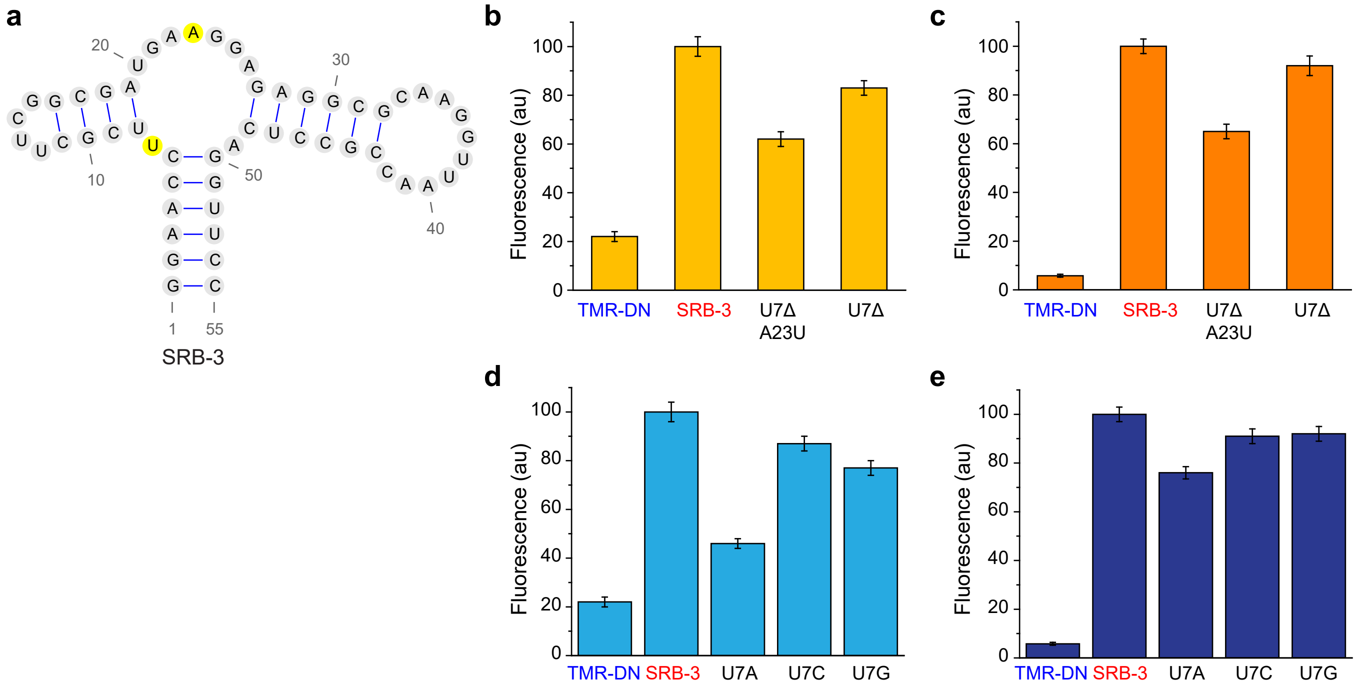


**Supplementary Figure 4. Mutations introduced in SRB-3 enhance its fluorescence. a)** Predicted secondary structure of SRB-3. **b,c)** Two mutations introduced into SRB-3 (highlighted in yellow) are beneficial for the fluorescence of SRB-3:TMR-DN, whereas two other modifications (mutation of $A_{23}$ to U and/or deletion of U7) have an adverse effect. For this comparison, SRB-3 at two different concentrations (7.5 nM in **b** and 750 nM in **c**) was mixed with TMR-DN (5 nM). At high SRB-3 concentration, the complexation is saturated and the measured fluorescence intensity is proportional to the quantum yield of the aptamer:TMR-DN complex. At low concentration, the fluorescence intensity depends on the quantum yield as well as the equilibrium dissociation coefficient of the complex. At both concentrations, SRB-3 performed better than the mutants. **d,e)** U is the preferred nucleotide at position 7 in SRB-3 as mutations to A, C or G decrease the fluorescence of SRB-3:TMR-DN. For these measurements, mutants at two different concentrations (7.5 nM in **d** and 750 nM in **e**) were mixed with TMR-DN (5 nM). In both cases, SRB-3 performed better than the tested mutants.

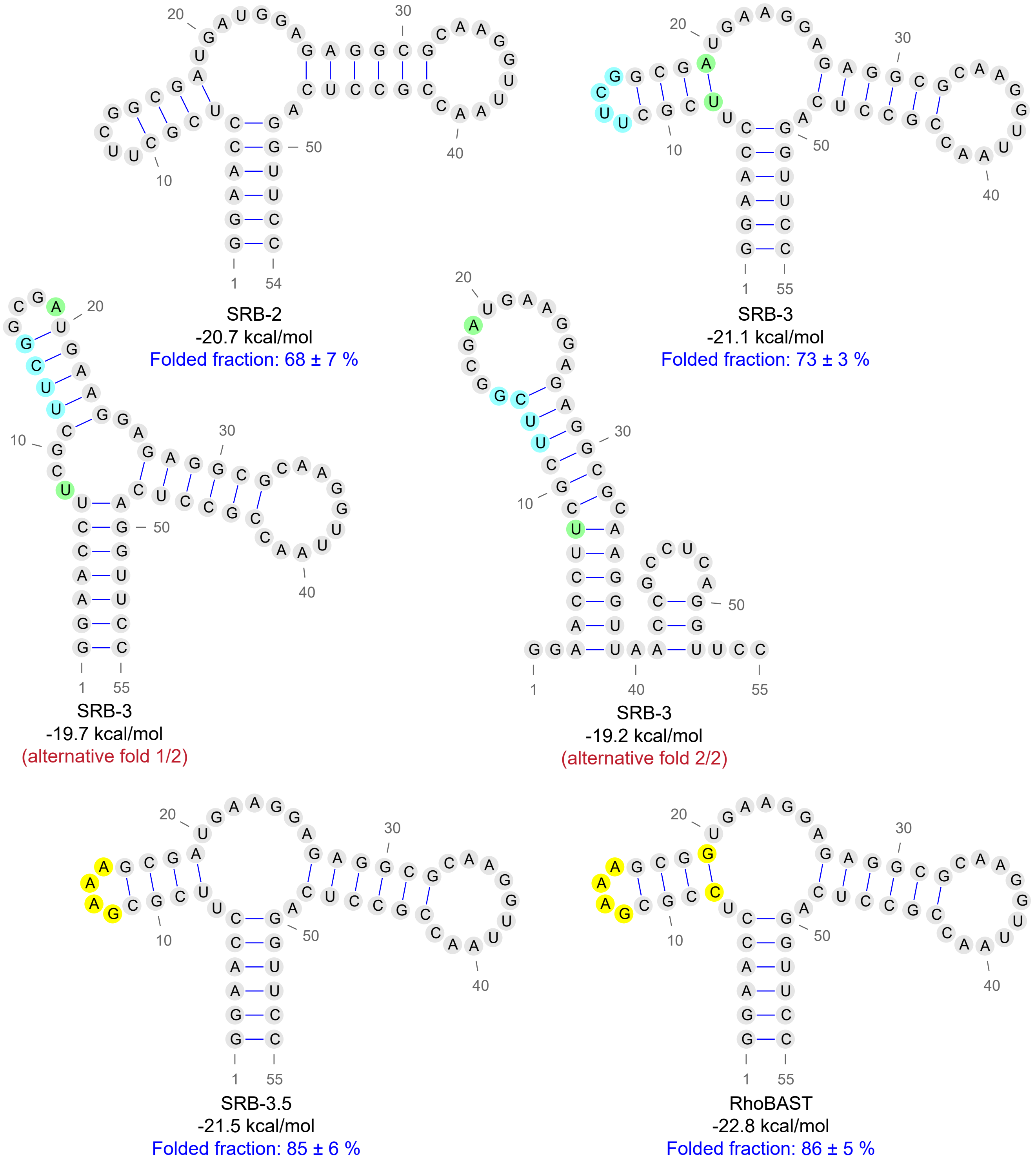


**Supplementary Figure 5. Alternative secondary structures of SRB-3 and the design of RhoBAST.** The alternative folds of SRB-3 were destabilized and thus eliminated by exchanging the UUCG tetraloop by GAAA and the $U_8$-$A_{19}$ base pair by C-G. The resulting RhoBAST sequence features a properly folded fraction of 86 ± 5%.

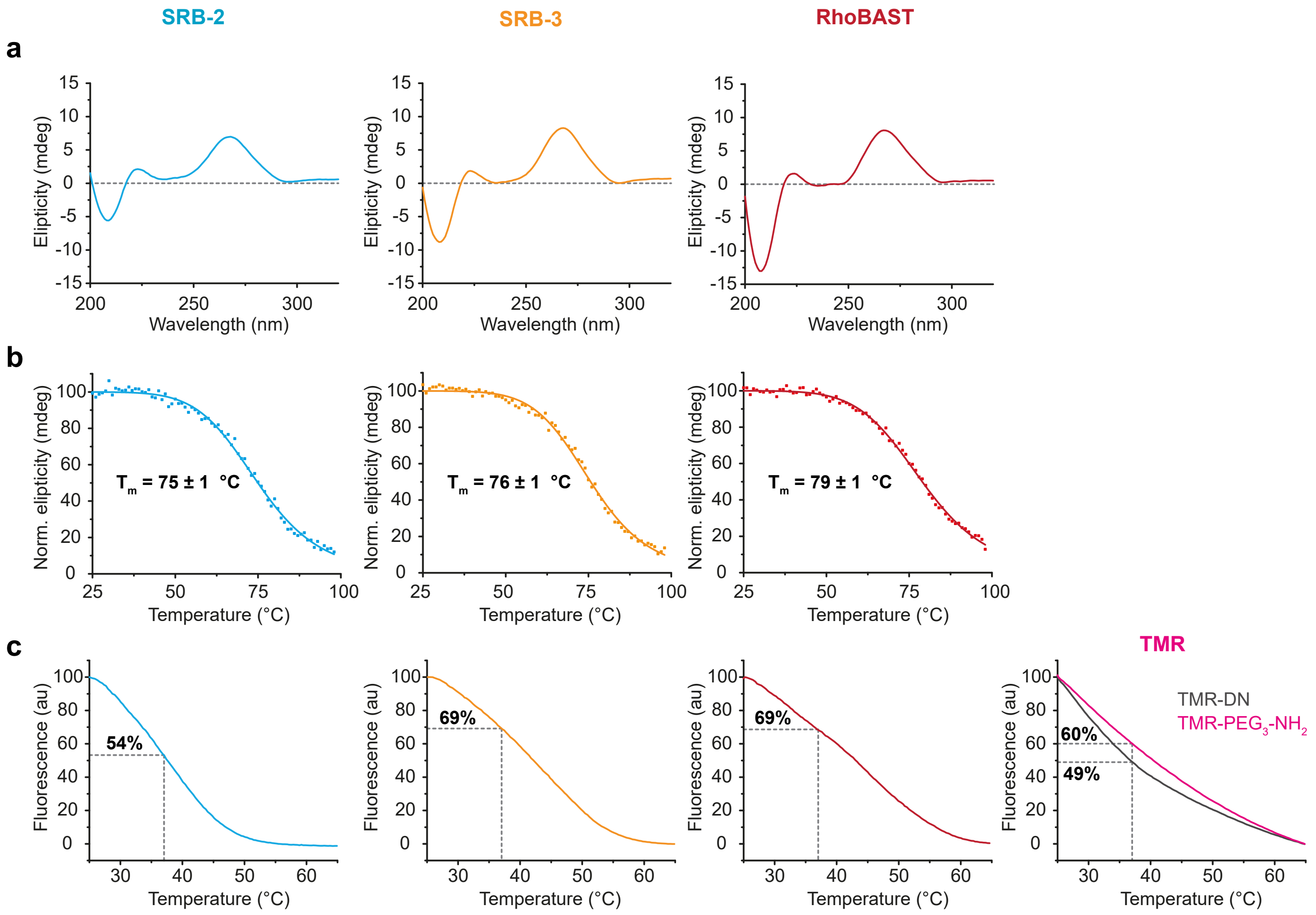


**Supplementary Figure 6. Characterization of SRB-2, SRB-3 and RhoBAST. a)** CD spectra at 25 °C. **b)** Melting curves, obtained by plotting the normalized ellipticity of the aptamers at 268 nm as a function of temperature from 25 °C to 98 °C. Shown are representative data points along with the best fit curve from fitting to the Boltzmann sigmoid. The melting temperature, $T_m$, represents the temperature at which the ellipticity decreases to 50% of its initial value at 25 °C. **c)** Temperature dependence of the fluorescence intensity of the aptamer:TMR-DN complexes and, for comparison, TMR-DN and TMR conjugated to the 2,2′-(ethylenedioxy)bis(ethylamine) linker, denoted as TMR-PEG$_3$-NH$_2$. Dashed lines indicate the fractional decreases in intensity upon heating from 25 to 37 °C.

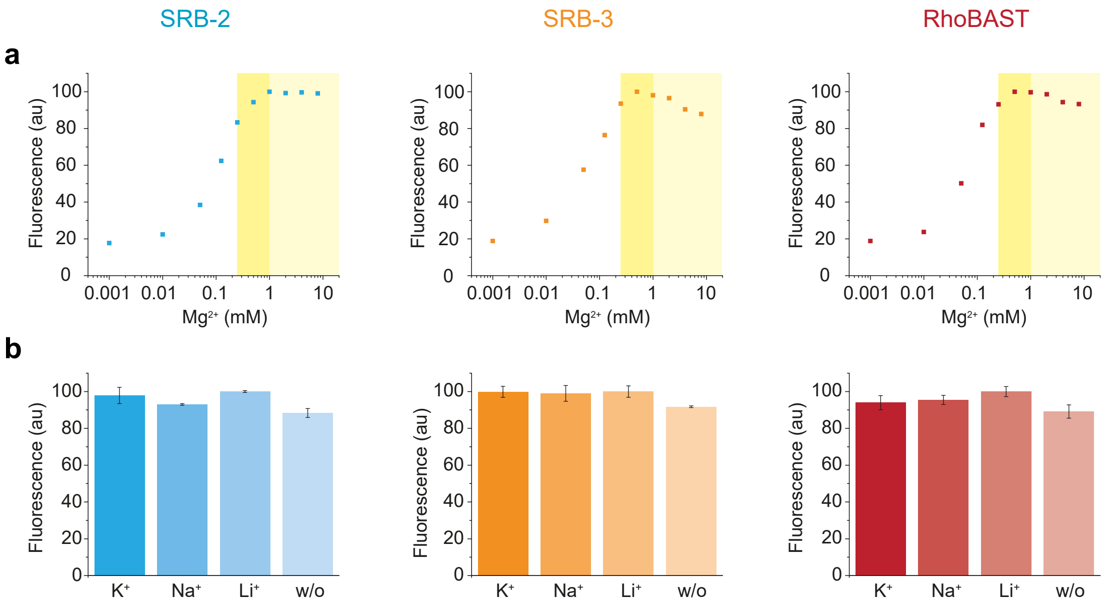


**Supplementary Figure 7. Dependence of the aptamer:TMR-DN fluorescence on the presence of metal ions in the solvent. a)** $Mg^{2+}$ ions are essential for aptamer:TMR-DN complex formation. The fluorescence of the aptamer:TMR-DN complexes (500 nM) was determined in a buffer containing 20 mM HEPES (pH 7.4), 125 mM KCl and various concentrations of $MgCl_2$ (1 µM – 10 mM) . The yellow regions designate the physiologically relevant concentration range of $Mg^{2+}$ ions in mammalian (~0.25 – 1 mM) and bacterial cells (~1 – 10 mM). **b)** Dependence of the aptamer:TMR-DN fluorescence on potassium, sodium, and lithium ions. The fluorescence of the aptamer:TMR-DN complexes (500 nM) was determined in a buffer containing 20 mM HEPES (pH 7.4), 1 mM $MgCl_2$ and 125 mM LiCl, NaCl or KCl. No significant dependence on LiCl, NaCl or KCl was observed. Even in the absence of any alkali metal ions, the complexes retain more than 90% of their maximum fluorescence. All experiments were carried out at 25 °C. Data represent the mean ± s.d. of three independent measurements.

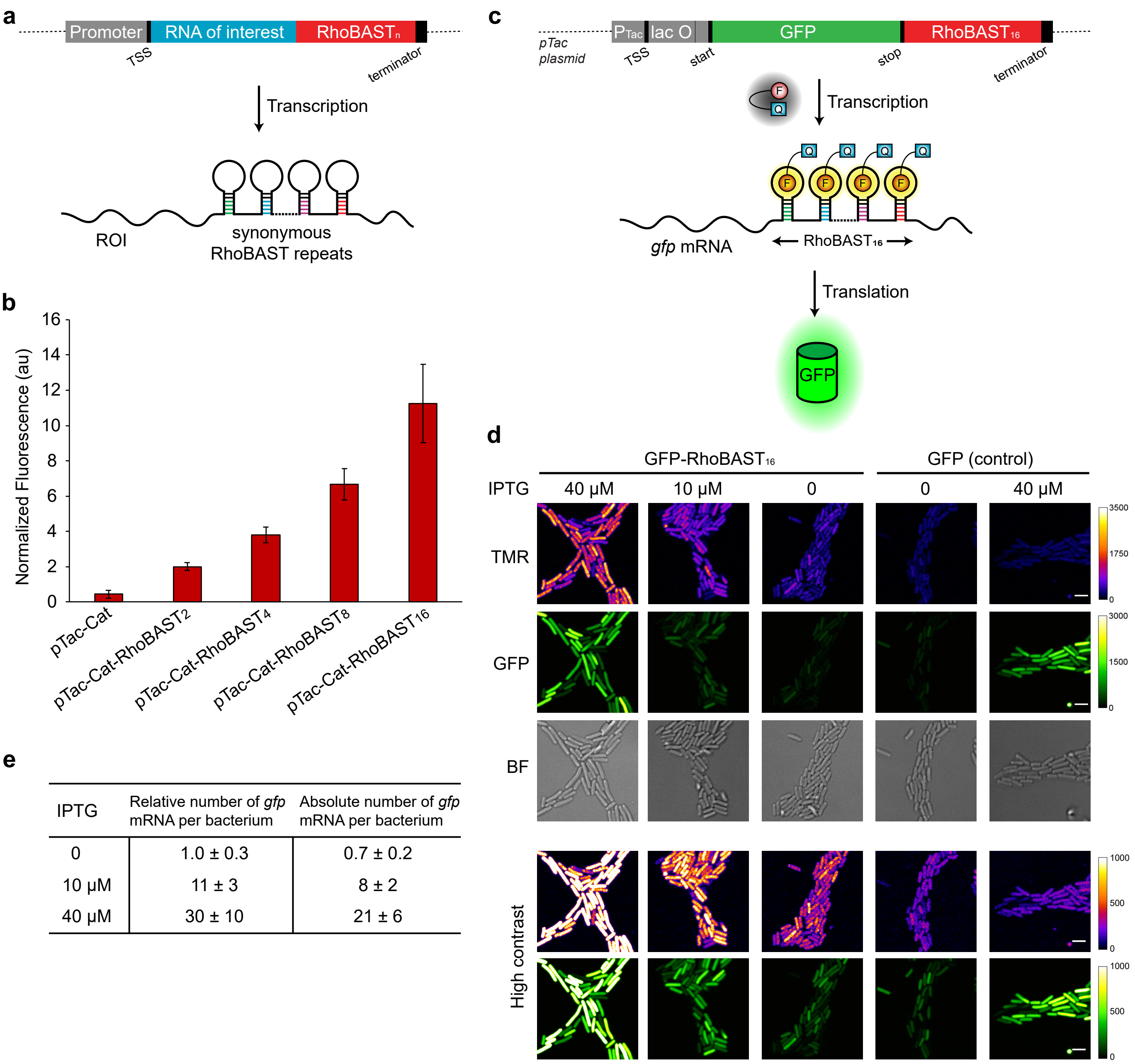


| IPTG | Relative number of *gfp* mRNA per bacterium | Absolute number of *gfp* mRNA per bacterium |
|---|---|---|
| 0 | 1.0 ± 0.3 | 0.7 ± 0.2 |
| 10 µM | 11 ± 3 | 8 ± 2 |
| 40 µM | 30 ± 10 | 21 ± 6 |

**Supplementary Figure 8. Synonymous repeats of RhoBAST increase the sensitivity of RNA imaging. a)** Scheme showing how to tag an RNA of interest (ROI) with synonymous repeats of RhoBAST at its 3′ end. Similarly, RhoBAST can be fused to an internal part or to the 5′ end of an ROI. ‘TSS’ denotes the transcription start site, ‘terminator’ indicates where the transcription ends. In order to promote correct folding of the aptamer repeats and to avoid homologous recombination between sequence repeats, synonymous repeats, different in sequence but identical in function, were generated mainly by using different terminal stem (H1) sequences. Sequences involved in H3 of RhoBAST can also be altered as long as they do not affect the affinity and the turn-on value of the aptamer. **b)** Dependence of the fluorescence intensity on the number of RhoBAST repeats. The fluorescence intensity (mean ± s.d) of fluorescence images of bacteria expressing *cat*, *cat-RhoBAST$_2$*, *cat-RhoBAST$_4$*, *cat-RhoBAST$_8$* or *cat-RhoBAST$_{16}$* mRNA reveals an almost linear fluorescence increase up to 16 repeats. The fluorescence intensity in bacteria expressing *cat-RhoBAST$_2$* is normalized to 2. **c)** IPTG-inducible plasmid system to express *gfp-RhoBAST$_{16}$* mRNA and GFP inside bacteria. RhoBAST$_{16}$ was fused to the 3′ UTR of GFP. **d)** Confocal images of bacteria expressing different amounts of *gfp-RhoBAST$_{16}$* mRNA. DH5α *E. coli* transformed with the pTac-GFP-RhoBAST$_{16}$ plasmid were grown at 37 °C, treated with different amounts of IPTG (0, 10, 40 µM), and fixed. Total RNA was also isolated from the same bacterial cultures in order to quantify the amount of *gfp-RhoBAST$_{16}$* mRNA at the time of fixation. Fixed bacteria were immobilized and imaged in M9 medium containing TMR-DN (20 nM). Bacteria transformed with pTac-GFP without the RhoBAST sequence were used as a control. Scale bars, 3 µm. **e)** Quantification of the average number of *gfp-RhoBAST$_{16}$* mRNA per bacterium. Total RNA was isolated from bacteria imaged in panel **d** and used in RT-qPCR for absolute quantification of RNA copy numbers. A small fragment (~180 nucleotides) of the GFP gene was *in-vitro* transcribed and used as a reference RNA in RT-qPCR.

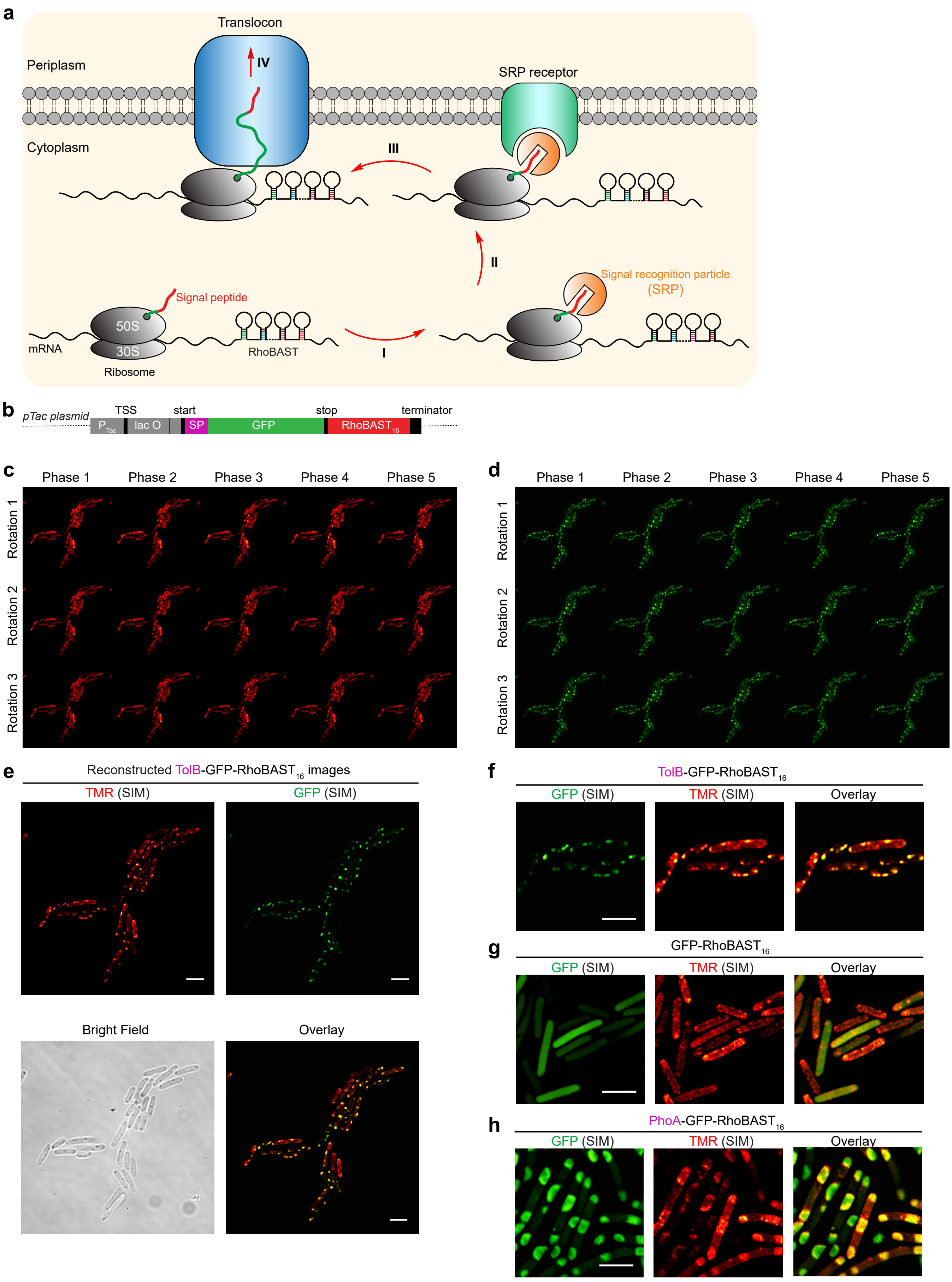
a
Translocon
IV
Periplasm
SRP receptor
Cytoplasm
III
II
Signal recognition particle
(SRP)
Signal peptide
50S
30S
mRNA
Ribosome
RhoBAST
I
b
pTac plasmid
TSS
start
stop
terminator
P Tac
lac O
SP
GFP
RhoBAST 16
c
Phase 1
Phase 2
Phase 3
Phase 4
Phase 5
Rotation 1
Rotation 2
Rotation 3
d
Phase 1
Phase 2
Phase 3
Phase 4
Phase 5
Rotation 1
Rotation 2
Rotation 3
e
Reconstructed TolB-GFP-RhoBAST16 images
TMR (SIM)
GFP (SIM)
Bright Field
Overlay
f
TolB-GFP-RhoBAST16
GFP (SIM)
TMR (SIM)
Overlay
g
GFP-RhoBAST16
GFP (SIM)
TMR (SIM)
Overlay
h
PhoA-GFP-RhoBAST16
GFP (SIM)
TMR (SIM)
Overlay

**Supplementary Figure 9. RhoBAST:TMR-DN imaging using SIM. a)** Scheme summarizing the basic principles of signal recognition particle (SRP) mediated cotranslational translocation of proteins. First, SRP binds to the signal sequence of the secretory protein while it emerges from the ribosome. Then, the entire SRP:mRNA:ribosome:nascent secretory protein chain ternary complex binds to the SRP receptor (FtsY) and the mRNA:ribosome:nascent secretory protein chain ternary complex is transferred to the translocase with the aid of the SRP receptor. Since the mRNA is tagged with RhoBAST, TMR fluorescence appears from the inner bacterial membrane. **b)** Plasmid system for bacterial expression of *gfp-RhoBAST*$_{16}$ mRNA containing a signal peptide (SP) sequence at the N-terminus of GFP. **c, d)** Raw SIM images of bacteria expressing *tolB-gfp-RhoBAST*$_{16}$ mRNA in the TMR (panel **c**) and the GFP (panel **d**) channel. TolB signal peptide binds to the SRP. **e)** Reconstructed SIM images of bacteria expressing *tolB-gfp-RhoBAST*$_{16}$ mRNA. **f)** Zoomed SIM images of bacteria expressing *tolB-gfp-RhoBAST*$_{16}$ mRNA. **g)** SIM images of bacteria expressing *gfp-RhoBAST*$_{16}$ mRNA. **h)** SIM images of bacteria expressing *phoA-gfp-RhoBAST*$_{16}$ mRNA. PhoA signal peptide binds SecB, but not SRP; therefore, the protein translocation occurs post-translationally. Scale bars, 3 µm.

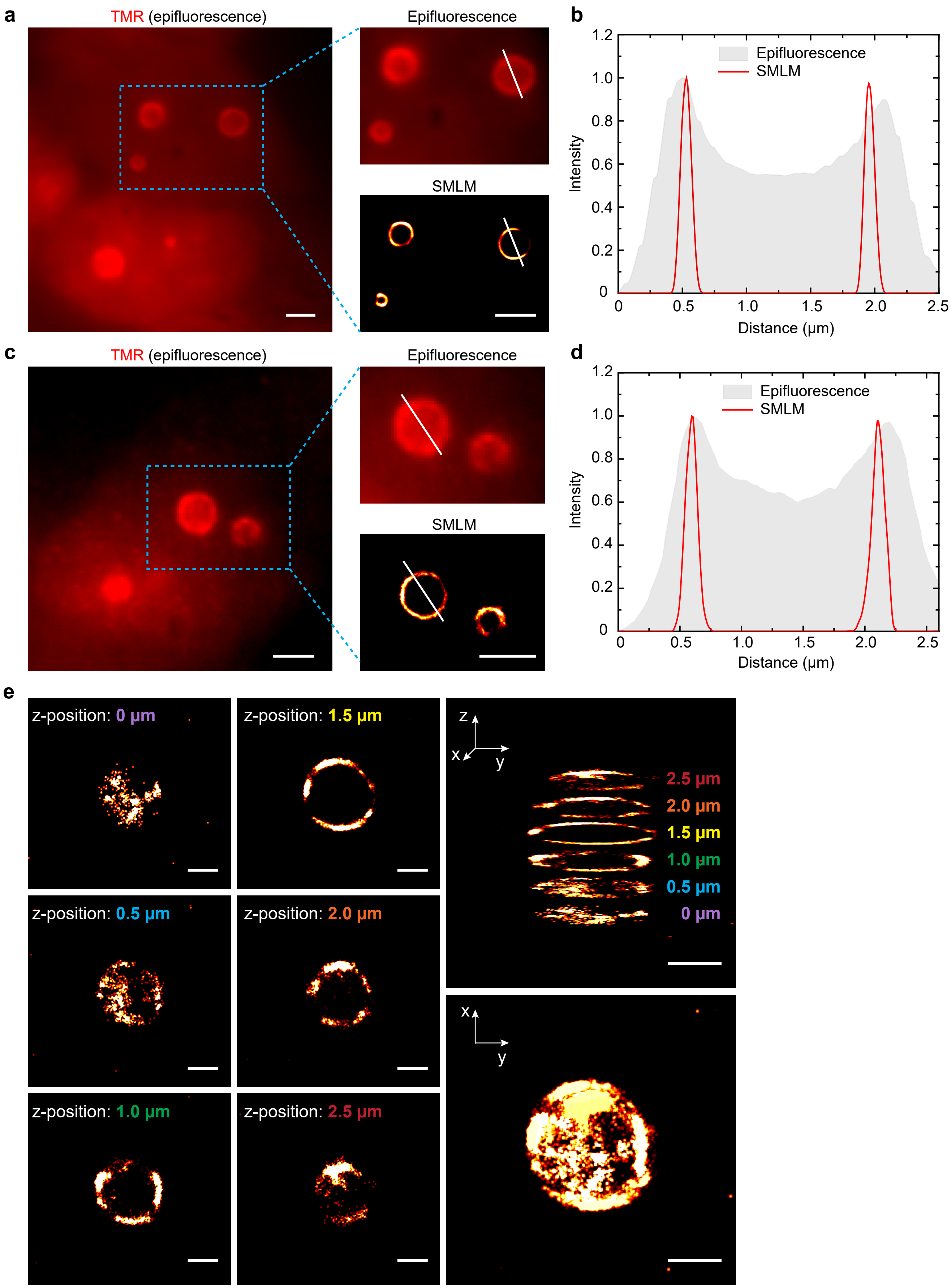
a
TMR (epifluorescence)
Epifluorescence
SMLM
b
Epifluorescence
SMLM
Intensity
Distance (μm)
c
TMR (epifluorescence)
Epifluorescence
SMLM
d
Epifluorescence
SMLM
Intensity
Distance (μm)
e
z-position: 0 μm
z-position: 1.5 μm
z-position: 0.5 μm
z-position: 2.0 μm
z-position: 1.0 μm
z-position: 2.5 μm
z
x
y
2.5 μm
2.0 μm
1.5 μm
1.0 μm
0.5 μm
0 μm

**Supplementary Figure 10. Epifluorescence and SMLM images of fixed HEK293T cells. a)** Epifluorescence and SMLM images of fixed HEK293T cells showing hollow spheres containing circular RhoBAST in the nuclei. Scale bars, 2 µm. **b)** Plot of the fluorescence intensity along the white lines in panel **a**, showing the much higher spatial resolution of SMLM. **c, d)** Same as panels **a** and **b**, showing different cells. **e)** SMLM images of a spherical subnuclear structure at different z-positions (left/middle), and a 3D representation (top right) and a maximum intensity projection (bottom right). Scale bars, 1 µm.

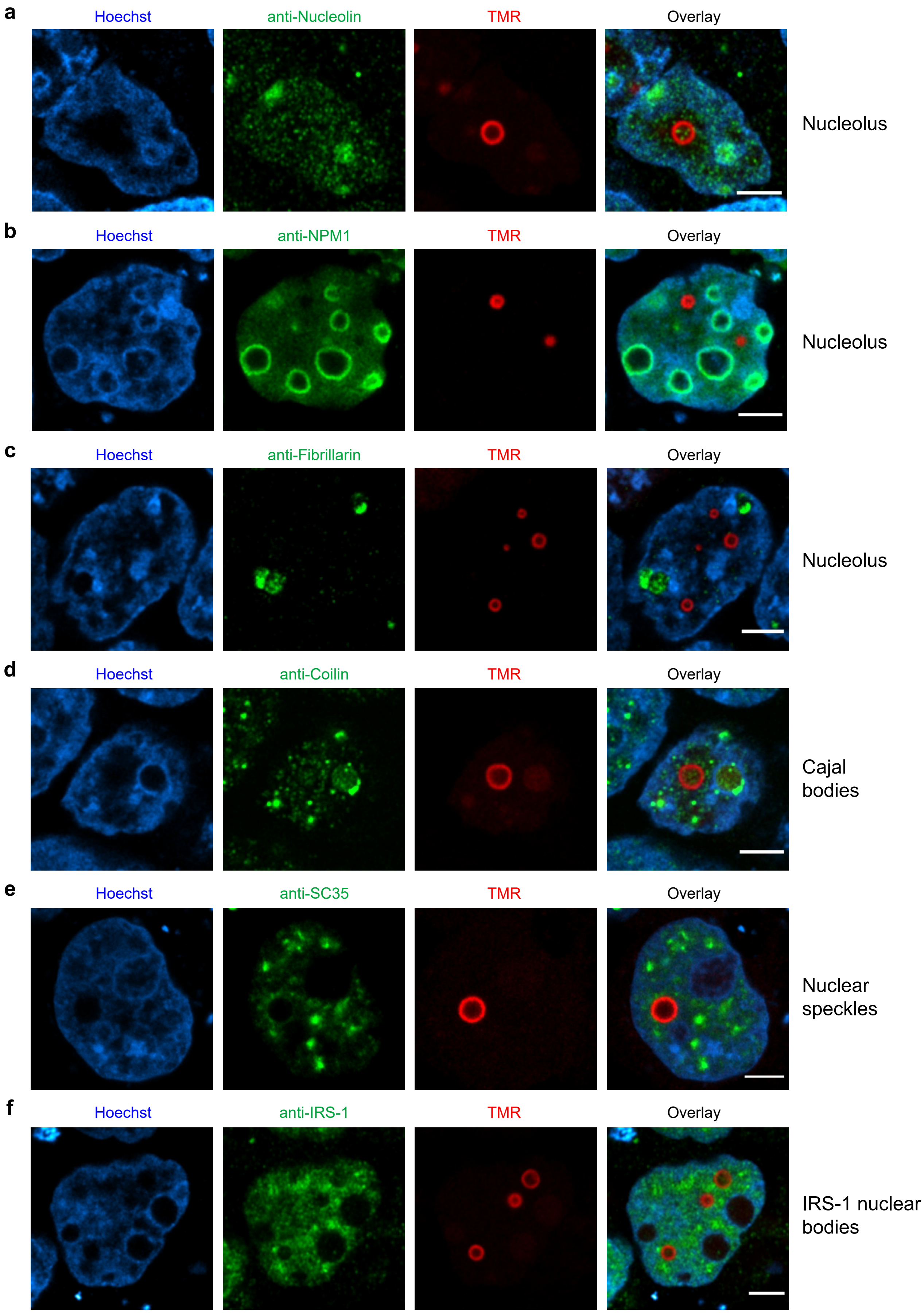
a
Hoechst
anti-Nucleolin
TMR
Overlay
Nucleolus
b
Hoechst
anti-NPM1
TMR
Overlay
Nucleolus
c
Hoechst
anti-Fibrillarin
TMR
Overlay
Nucleolus
d
Hoechst
anti-Coilin
TMR
Overlay
Cajal bodies
e
Hoechst
anti-SC35
TMR
Overlay
Nuclear speckles
f
Hoechst
anti-IRS-1
TMR
Overlay
IRS-1 nuclear bodies

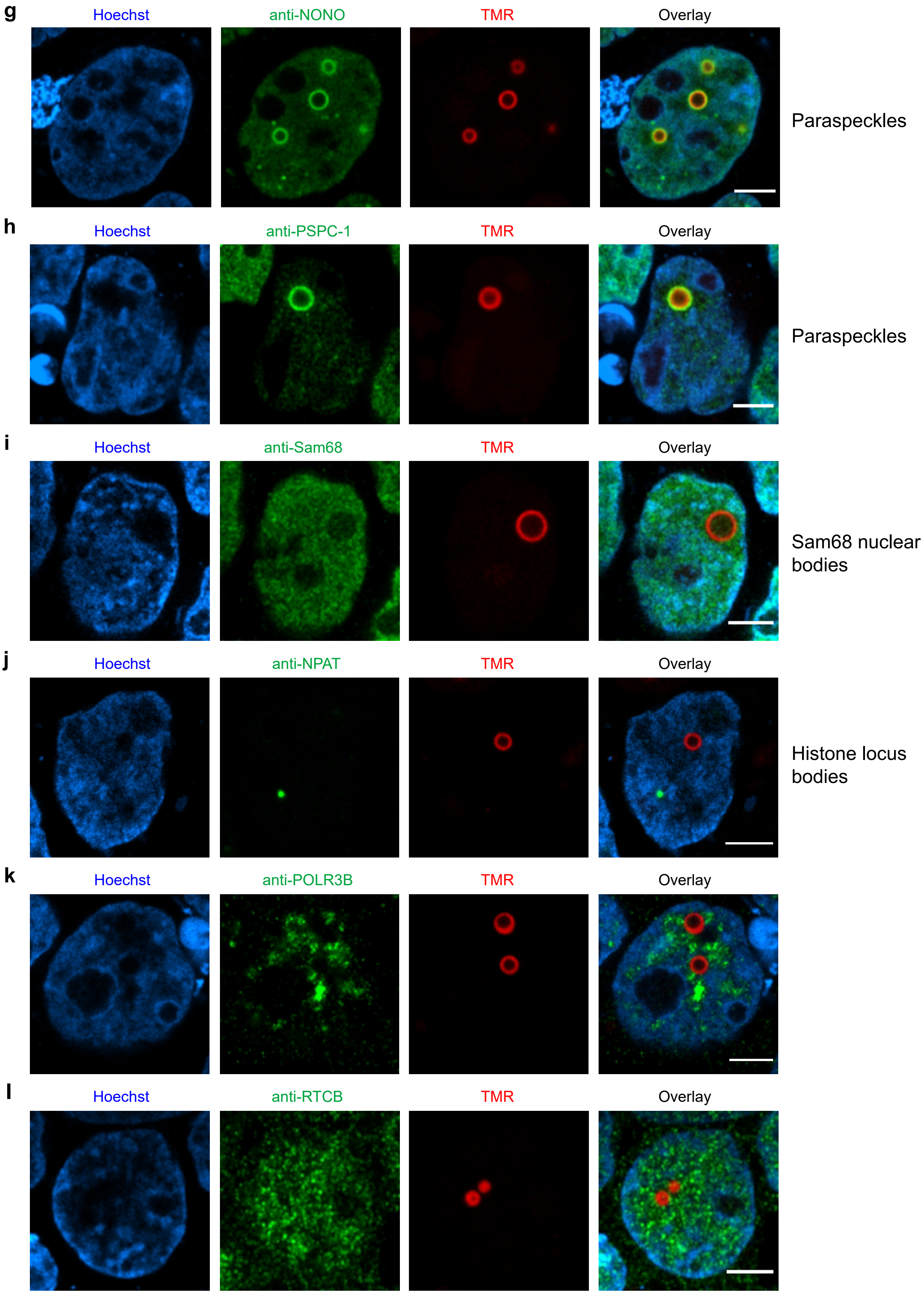
g
Hoechst
anti-NONO
TMR
Overlay
Paraspeckles
h
Hoechst
anti-PSPC-1
TMR
Overlay
Paraspeckles
i
Hoechst
anti-Sam68
TMR
Overlay
Sam68 nuclear bodies
j
Hoechst
anti-NPAT
TMR
Overlay
Histone locus bodies
k
Hoechst
anti-POLR3B
TMR
Overlay
l
Hoechst
anti-RTCB
TMR
Overlay

**Supplementary Figure 11. Confocal microscopy of subnuclear structures formed by circular RhoBAST reveals colocalization with paraspeckle proteins NONO and PSPC1**. Tornado-RhoBAST expressing HEK293T cells were fixed and stained with a primary antibody targeting common nuclear bodies, followed by an Alexa488-conjugated secondary antibody. Images were taken in the presence of TMR-DN (100 nM, red) and Hoechst (1 µg/mL, blue) in DPBS containing 1 mM $MgCl_2$. The following antibodies were used for this study: **a)** anti-Nucleolin (for nucleolus), **b)** anti-NPM1 (for nucleolus), **c)** anti-Fibrillarin (for nucleolus), **d)** anti-Coilin (for Cajal bodies), **e)** anti-SC35 (for nuclear speckles), **f)** anti-IRS1 (IRS-1 nuclear bodies), **g)** anti-NONO (for paraspeckles), **h)** anti-PSPC1 (for paraspeckles), **i)** anti-Sam68 (for Sam68 nuclear bodies), **j)** anti-NPAT (for histone locus bodies). In addition, **k)** anti-POLR3B and **l)** anti-RTCB were used because the former takes part in the transcription of Tornado-RhoBAST and the latter is responsible for circularization of the final transcript. Scale bars, 5 µm.

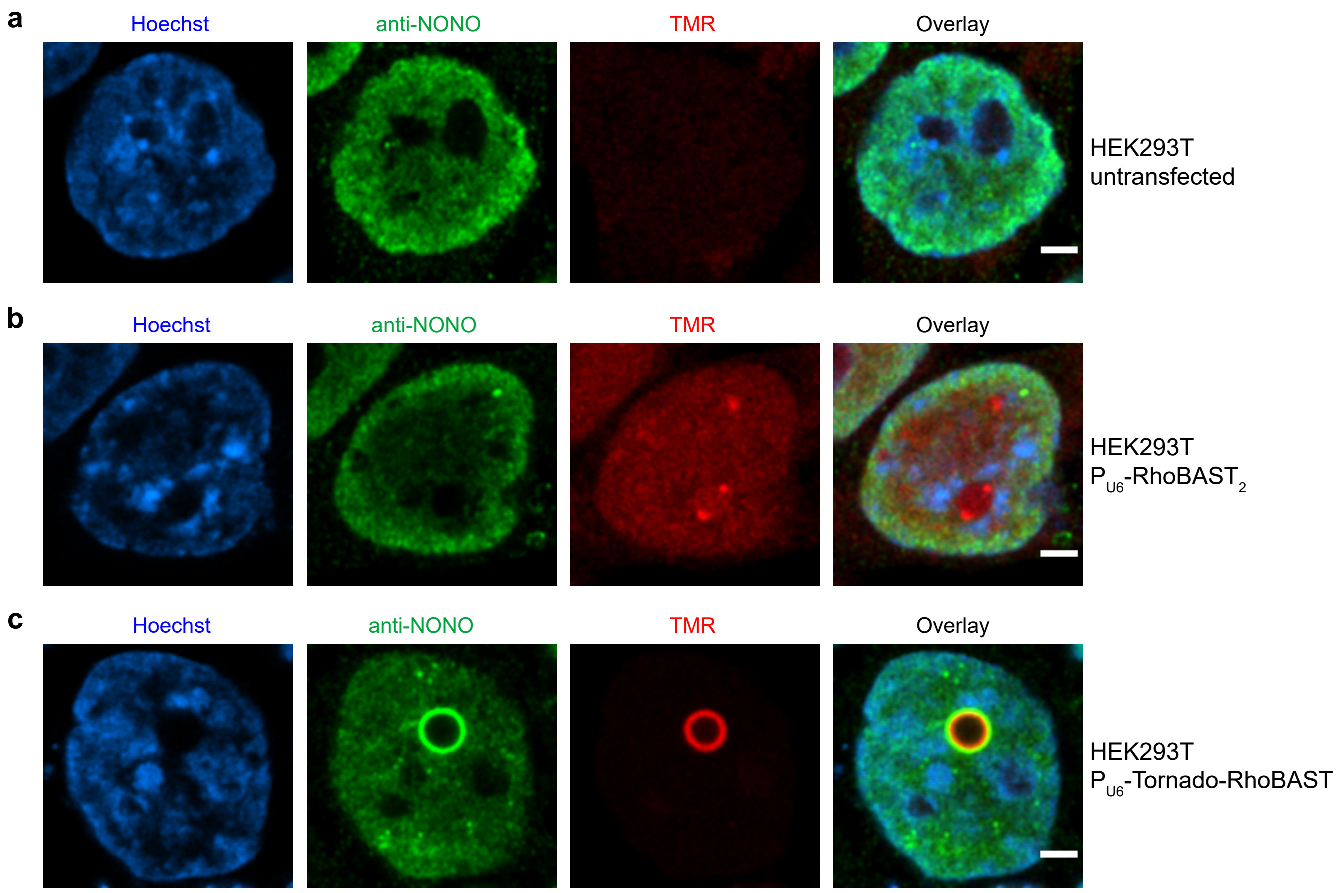


**Supplementary Figure 12. Hollow, spherical subnuclear structures are not observed in untransfected or linear-RhoBAST$_2$ expressing cells. a)** Untransfected, **b)** P$_{U6}$-RhoBAST$_2$ and **c)** P$_{U6}$-Tornado-RhoBAST expressing HEK293T cells were fixed and stained with the anti-NONO primary antibody followed by an Alexa488-conjugated secondary antibody. Confocal images were taken in the presence of TMR-DN (100 nM, red) and Hoechst (1 µg/mL, blue) in DPBS containing 1 mM $MgCl_2$. To avoid saturation of the pixels in the TMR channel of panel **c**, ~10-fold less laser power (561 nm) was used compared to panels **a** and **b**. Hollow, spherical subnuclear structures were only observed in Tornado-RhoBAST expressing cells, suggesting that the high concentration of cyclic RhoBAST in the nucleus drives formation of these structures. Scale bars, 3 µm.

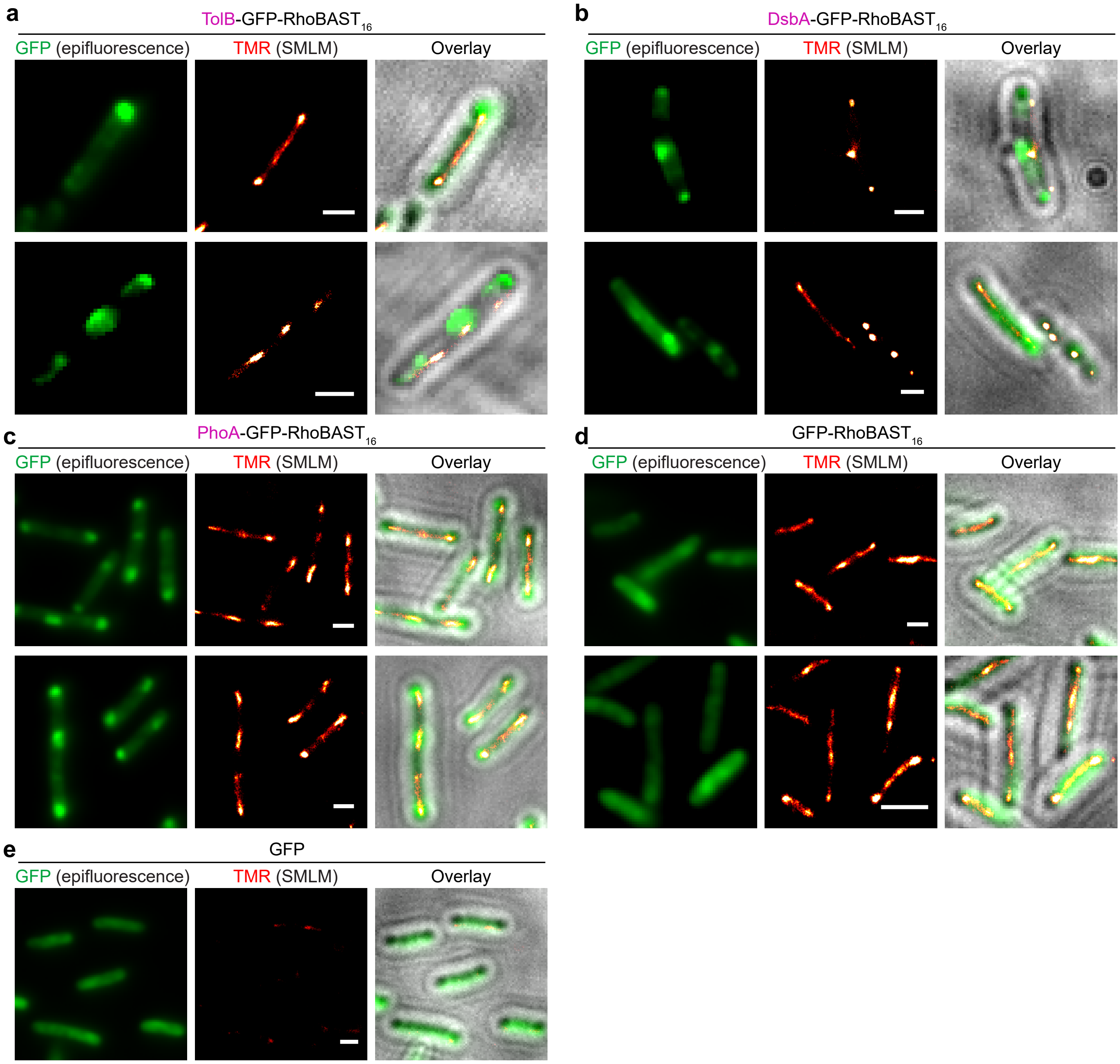


**Supplementary Figure 13. SMLM images of fixed *E. coli* bacteria expressing *gfp-RhoBAST$_{16}$* fused to different signal peptides. a) – e)** Epifluorescence images of GFP fluorescence (left), SMLM images of TMR-DN (middle) and overlay of both with the bright field image (right) for **a)** *tolB-gfp-RhoBAST$_{16}$*, **b)** *dsbA-gfp-RhoBAST$_{16}$*, **c)** *phoA-gfp-RhoBAST$_{16}$*, **d)** *gfp-RhoBAST$_{16}$* and **e)** *gfp* mRNA. The DsbA signal peptide binds SRP; therefore, the protein translocation occurs co-translationally. Scale bars, 1 µm. Average localization accuracies of the SMLM images were found to be in the range 6 – 30 nm.

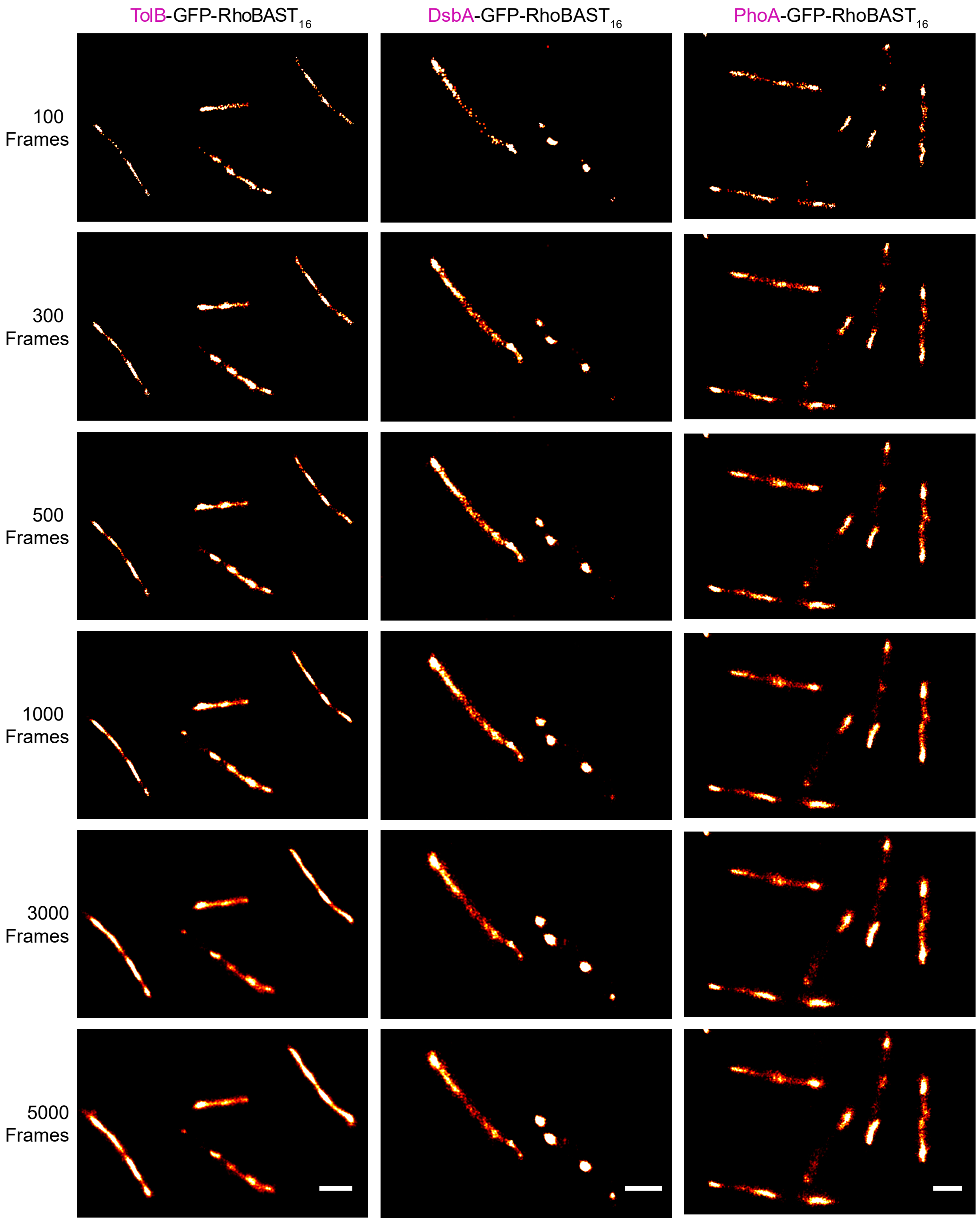


**Supplementary Figure 14. Comparison of SMLM images of fixed *E. coli* bacteria reconstructed with different numbers of camera frames.** SMLM images are shown for TMR-DN bound to *tolB-gfp-RhoBAST$_{16}$* (left), *dsbA-gfp-RhoBAST$_{16}$* (middle) and *phoA-gfp-RhoBAST$_{16}$* mRNAs (right), reconstructed for 100 – 5,000 frames (100 ms each). Scale bars, 1 µm. Image intensities were normalized to the maximum intensity value in each image.

**Supplementary Table 1.** Photophysical and binding properties of different FLAPs.

| FLAP | $\lambda_{exc}$ (nm) | $\lambda_{em}$ (nm) | $\varepsilon$ ($M^{-1}cm^{-1}$) | QY | Relative Brightness | Turn-on | $K_D$ (nM) | $k_a$ ($M^{-1}s^{-1}$) | $k_d$ ($s^{-1}$) |
|---|---|---|---|---|---|---|---|---|---|
| RhoBAST:TMR-DN | 564 | 590 | 96000 | 0.57 ± 0.04 | 460 | 26 | 15 ± 1 | $1.8 \times 10^7$ | 0.66 |
| SRB-3:TMR-DN | 564 | 590 | 94000 | 0.56 ± 0.03 | 440 | 25 | 20 ± 1 | - | - |
| SRB-2:TMR-DN [1] | 564 | 590 | 90500 | 0.33 ± 0.04 | 250 | 17 | 35 ± 1 | - | - |
| SRB-2:SR-DN [2] | 579 | 596 | 85200 | 0.65 | 470 | 105 | 1340 | - | - |
| o-Coral:Gemini-561 [3] | 580 | 596 | 70500 [a] | 0.58 | 350 | 13 | 73 | - | - |
| SiRA:SiR-$PEG_3$-$NH_2$ [4] | 649 | 662 | 86000 | 0.98 | 710 | 7 | 430 | $6.6 \times 10^3$ | 0.0028 |
| DNB:SR-DN [5] | 572 | 591 | 50250 | 0.98 | 420 | 56 | 800 | - | - |
| Riboglow:Cbl-Atto590 [6] | 594 | 622 | 120000 | 0.31 | 310 | 5 | 34 | - | - |
| Broccoli-DFHBI-1T [7] | 470 | 505 | 28900 | 0.41 | 100 | 1000 | 305 | $1.5 \times 10^4$ | 0.0036 |
| Broccoli–DFHBI-1T [8] | - | - | - | - | - | - | - | $5.4 \times 10^4$ | 0.018 |
| Broccoli-BI [7] | 470 | 505 | 33600 | 0.67 | 190 | 1390 | 51 | $1.4 \times 10^4$ | 0.0006 |
| Corn-DFHO [8] | 505 | 545 | 29000 | 0.25 | 60 | 420 | 70 | $2.3 \times 10^4$ | 0.008 |
| Mango TO1-Biotin [9] | 510 | 535 | 77500 | 0.14 | 90 | 1100 | 3.9 | - | - |
| Mango IV [10] | 510 | 535 | 76000 | 0.42 | 270 | 3240 | 11 ± 1 | - | - |
| Pepper530 [11] | 485 | 530 | 65300 | 0.66 | 360 | 3600 | 3.5 | $6.6 \times 10^5$ | 0.0023 |
| Pepper620 [11] | 577 | 620 | 100000 | 0.58 | 490 | 12600 | 6.1 | - | - |

[a] The reported extinction coefficient was divided by two since the system is dimeric.

**Supplementary Table 2.** Association ($k_a$) and dissociation ($k_d$) rate coefficients, and average time intervals ($T$) between successive dye binding events for different dye concentrations ($c$).[a]

| FLAP | $k_a$ ($M^{-1}s^{-1}$) | $k_d$ ($s^{-1}$) | $T$ (s) ($c$ = 1 nM) | $T$ (s) ($c$ = 10 nM) | $T$ (s) ($c$ = 100 nM) | $T$ (s) ($c$ = 1 µM) |
|---|---|---|---|---|---|---|
| RhoBAST:TMR-DN | $1.8 \times 10^7$ | 0.66 | $5.7 \times 10^1$ | 7.1 | 2.1 | 1.6 |
| SiRA:SiR-PEG$_3$-NH$_2$ | $6.6 \times 10^3$ | 0.0028 | $1.5 \times 10^5$ | $1.6 \times 10^4$ | $1.9 \times 10^3$ | $5.1 \times 10^2$ |
| Broccoli-DFHBI-1T | $1.5 \times 10^4$ | 0.0036 | $6.7 \times 10^4$ | $6.9 \times 10^3$ | $9.4 \times 10^2$ | $3.4 \times 10^2$ |
| Broccoli-BI | $1.4 \times 10^4$ | 0.0006 | $7.3 \times 10^4$ | $8.8 \times 10^3$ | $2.4 \times 10^3$ | $1.7 \times 10^3$ |
| Corn-DFHO | $2.3 \times 10^4$ | 0.008 | $4.4 \times 10^4$ | $4.5 \times 10^3$ | $5.6 \times 10^2$ | $1.7 \times 10^2$ |
| Pepper530 | $6.6 \times 10^5$ | 0.0023 | $2.0 \times 10^3$ | $5.9 \times 10^2$ | $4.5 \times 10^2$ | $4.4 \times 10^2$ |

[a] $T$ was calculated as $\tau_{ON} + \tau_{OFF} = (k_d)^{-1} + (c \times k_a)^{-1}$.

**Supplementary Table 3.** DNA template sequences of the aptamers used in this study

| Name | DNA Sequence |
|---|---|
| SRB-2 | T7 φ6.5-GGAACCTCGCTTCGGCGATGATGGAGAGGCGCAAGGTTAACCGCCTCAGGTTCC |
| SRB-3 | T7 φ6.5-GGAACCTTCGCTTCGGCGATGAAGGAGAGGCGCAAGGTTAACCGCCTCAGGTTCC |
| RhoBAST | T7 φ6.5-GGAACCTCCGCGAAAGCGGTGAAGGAGAGGCGCAAGGTTAACCGCCTCAGGTTCC |
| RhoBAST$_2$ | AGAGTCGACATA<br>GAGGAACCTCCGCGAAAGCGGTGAAGGAGAGGCGCAAGGTTAACCGCCTCAGGTTCCTC ATA<br>ACAAGGCCTCCGCGAAAGCGGTGAAGGAGCGGCACAAGGTTAACTGCCGCAGGCCTTGT<br>ATACTCGAGAGA |
| RhoBAST$_4$ | AGAGTCGACATA<br>GAGGAACCTCCGCGAAAGCGGTGAAGGAGAGGCGCAAGGTTAACCGCCTCAGGTTCCTC ATA<br>ACAAGGCCTCCGCGAAAGCGGTGAAGGAGCGGCACAAGGTTAACTGCCGCAGGCCTTGT ATA<br>CTCGACATA<br>GGAAGACCTCCGCGAAAGCGGTGAAGGAGCGGTGCAAGGTTAACCACCGCAGGTCTTCC ATA<br>AGCAGACCTCCGCGAAAGCGGTGAAGGAGTGGCGCAAGGTTAACCGCCACAGGTCTGCT ATA<br>CTCGAGAGA |
| Tornado-<br>RhoBAST | GTCGACGGGCCGCACTCGCCGGTCCCAAGCCCGGATAAAATGGGAGGGGGCGGGAAACCGCCTA<br>ACCATGCCGAGTGCGGCCGCACCTCCGCGAAAGCGGTGAAGGAGAGGCGCAAGGTTAACCGCCT<br>CAGGTGTGGCCGCGGTCGGCGTGGACTGTAGAACACTGCCAATGCCGGTCCCAAGCCCGGATAA<br>AAGTGGAGGGTACAGTCCACGCTCTAGA |
| Biotin-<br>RhoBAST-Cy5 | T7 φ2.5–AGGAACCTCCGCGAAAGCGGTGAAGGAGAGGCGCAAGGTTAACCGCCTCAGGTTCC<br>AA |
| T7 φ6.5 | TAATACGACTCACTATA (for transcripts starting with G) |
| T7 φ2.5 | TAATACGACTCACTATT (for transcripts starting with A) |

**Supplementary Table 4.** Sequences of the DNA cassettes used for the genome modification of *E. coli*

| | |
|---|---|
| **OmpA cassette** | 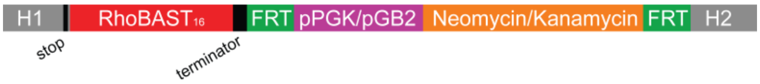  |
| Homologous Arm 1 (H1) | TCGTCGCGTAGAGATCGAAGTTAAAGGTATCAAAGACGTTG**TAA** |
| Homologous Arm 2 (H2) | CCTTACCCAGCAATGCCTGCAGATCCTGCTTCAGAGAAGA |
| Terminator | GTTCTCGTCTGGTAGAAAAACCCCGCTGCTGCGGGGTTTTTTTTGCCTTTAGTAAATTGA |
| **RNaseE cassette** | 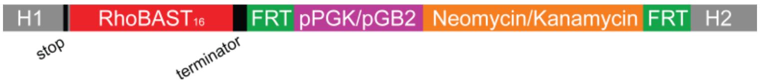  |
| Homologous Arm 1 (H1) | GCAGGTGGTCATACGGCAACACATCATGCCTCTGCCGCTCCTGCGCGTCCGCAACCTGTTGAG**TAA** |
| Homologous Arm 2 (H2) | ACATATTAAA ATTCATCGAA TGGCATCCTT GCTAACCAAC AATGCAAAAT AGGCAA |
| Terminator | TAATTAGCTCAAAGTAATCAAGCCCTGGTAACTGCCAGGGCTTTTTTATTTCATCTTTGAATCA |
| **Cat cassette** | 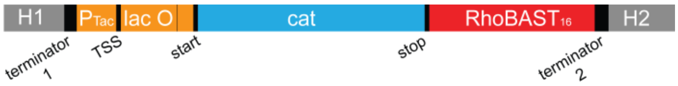  |
| Homologous Arm 1 (H1) | ATCATCCTCTGCATGGTCAGGTCATGGATGAGCAGACGATGGTGCAGGAT |
| Homologous Arm 2 (H2) | ATCCTGCTGATGAAGCAGAACAACTTTAACGCCGTGCGCTGTTCGCATTATCCGAACCAT |
| Terminator 1 | ACAGCGAAAAAACCCCGCCCTGTCAGGGGCGGGGTTTTTTGCGCGT |

**Supplementary Table 5.** Summary of antibodies used in this study.

| **Antibody** | **Source** | **Company** | **Catalog Number** | **Dilution Factor** |
|---|---|---|---|---|
| Anti-Nucleolin | Rabbit | Thermo Fisher | PA5-82860 | 1:50 |
| Anti-NPM1 | Mouse | Thermo Fisher | 32-5200 | 1:250 |
| Anti-Fibrillarin | Mouse | Thermo Fisher | 480009 | 1:250 |
| Anti-RTCB | Rabbit | Thermo Fisher | PA5-64867 | 1:25 |
| Anti-NONO | Rabbit | Sigma Aldrich | N8789 | 1:200 |
| Anti-PSPC1 | Rabbit | Thermo Fisher | PA5-58585 | 1:50 |
| Anti-Sam68 | Rabbit | Sigma Aldrich | HPA051280 | 1:100 |
| Anti-NPAT | Rabbit | Thermo Fisher | PA5-66839 | 1:100 |
| Anti-POLR3B | Rabbit | Thermo Fisher | PA5-57671 | 1:50 |
| Anti-Coilin | Mouse | Sigma Aldrich | C1862 | 1:500 |
| Anti-SC35 | Mouse | Sigma Aldrich | S4045 | 1:1000 |
| Anti-IRS1 | Rabbit | Thermo Fisher | PA1-1057 | 1:250 |
| Anti-Mouse IgG Alexa488 | Goat | Thermo Fisher | A28175 | 1:1000 |
| Anti-Rabbit IgG Alexa488 | Goat | Thermo Fisher | A11008 | 1:500 |

**Supplementary Note:** Synthesis of TMR-SS-Biotin and TMR-DN

TMR-SS-Biotin was synthesized as described in the scheme below:

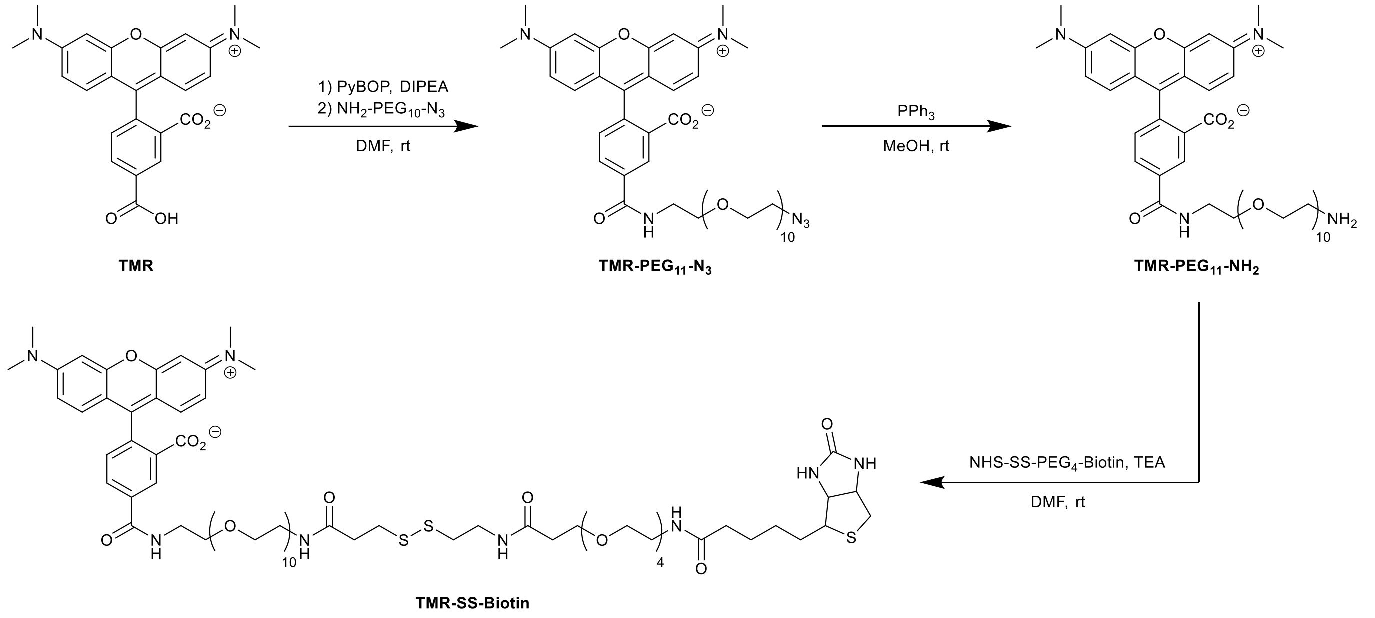


### a) Synthesis of TMR-PEG$_{11}$-N$_3$

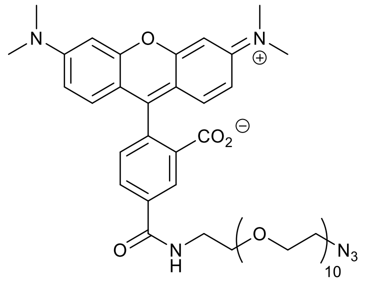

$C_{47}H_{66}N_6O_{14}$
939.07 g/mol

PyBOP (62.9 mg, 121 µmol, 1.20 eq) and 5-carboxytetramethylrhodamine (43.3 mg, 101 µmol, 1.00 eq) were dissolved in 6 mL dry DMF. Then, DIPEA (20.5 µL, 121 µmol, 1.20 eq) was added and the reaction mixture was stirred for 2 h at room temperature. O-(2-Aminoethyl)-O′-(2-azidoethyl) nonaethylene glycol (53.0 mg, 101 µmol, 1.00 eq) was dissolved in 1 mL dry DMF and added into the mixture. The reaction was stirred overnight at room temperature and the solvent was removed under reduced pressure. The crude mixture was purified by silica column chromatography ($CHCl_3$:MeOH:$H_2O$ = 20:1:0 → 10:1:0.1) and the product was obtained as dark violet solid (46.0 mg, 49.0 µmol, 49%).

$^{1}$**H NMR** ($CD_3OD$, 500 MHz): $\delta$ = 3.26 (s, 12 H), 3.36 (t, $J$=5.0 Hz, 2 H), 3.59 - 3.70 (m, 40 H), 3.72 (d, $J$=5.3 Hz, 2 H), 6.89 (d, $J$=2.1 Hz, 2 H), 7.00 (dd, $J$=9.5, 2.1 Hz, 2 H), 7.24 (d, $J$=9.4 Hz, 2 H), 7.36 (d, $J$=7.9 Hz, 1 H), 8.07 (dd, $J$=7.9, 1.8 Hz, 1 H), 8.55 (d, $J$=1.7 Hz, 1 H) ppm.

$^{13}$**C{$^{1}$H} NMR** (126 MHz, $CD_3OD$): $\delta$= 41.0, 41.3, 51.9, 70.7, 71.3, 71.6, 71.7, 71.7, 71.8, 71.8, 97.5, 114.8, 115.1, 129.7, 129.7, 130.9, 132.7, 137.3, 137.5, 141.9, 158.7, 159.1, 160.7, 169.3, 172.2 ppm.

**MS** (HR-ESI, pos): measured m/z = 961.4531, calculated m/z = 961.4529 for $C_{47}H_{66}N_6NaO_{14}$ $[M+Na]^+$.

**b) Synthesis of TMR-PEG$_{11}$-NH$_2$**

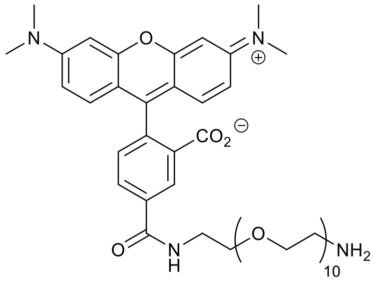


$C_{47}H_{68}N_4O_{14}$
913.08 g/mol

TMR-PEG$_{11}$-N$_3$ (86.0 mg, 91.6 µmol, 1.00 eq) was dissolved in 10 mL MeOH and triphenylphosphine (72.1 mg, 275 µmol, 3.00 eq) was added. The mixture was stirred overnight at room temperature. Next, 1 mL water was added and the solution was stirred for an additional 10 min. The solvent was removed under reduced pressure and the crude mixture was purified by silica column chromatography ($CHCl_3$:MeOH:$NH_3$(28% in $H_2O$) = 10:1:0 → 6:1:0.1) and the product was obtained as dark violet solid (65.1 mg, 71.2 µmol, 78%).

$^{1}$**H NMR** ($CD_3OD$, 500 MHz): $\delta$ = 2.91 (t, $J$=5.3 Hz, 2 H), 3.28 (s, 12 H), 3.58 - 3.70 (m, 40 H), 3.72 (t, $J$=5.3 Hz, 2 H), 6.91 (d, $J$=2.6 Hz, 2 H), 7.01 (dd, $J$=9.5, 2.6 Hz, 2 H), 7.24 (d, $J$=9.4 Hz, 2 H), 7.36 (d, $J$=7.9 Hz, 1 H), 8.06 (dd, $J$=7.9, 1.8 Hz, 1 H), 8.53 (d, $J$=1.8 Hz, 1 H) ppm.

$^{13}$**C{$^{1}$H} NMR** (126 MHz, $CD_3OD$): $\delta$= 41.0, 41.3, 41.8, 70.7, 71.2, 71.5, 71.6, 71.7, 71.8, 97.5, 114.9, 115.1, 129.6, 129.7, 130.9, 132.7, 137.3, 137.3, 142.2, 158.8, 159.1, 161.2, 169.3, 172.4 ppm.

**MS** (HR-ESI, pos): measured m/z = 913.4815, calculated m/z = 913.4805 for $C_{47}H_{69}N_4O_{14}$ $[M+H]^+$.

**c) Synthesis of TMR-SS-Biotin**

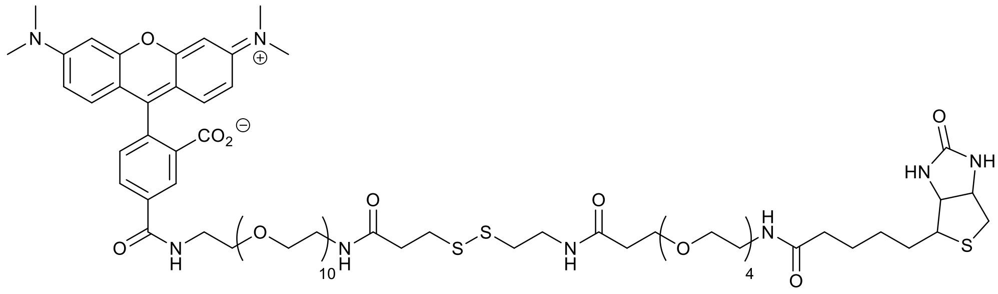

$C_{73}H_{112}N_8O_{22}S_3$
1549.91 g/mol

TMR-PEG$_{11}$-NH$_2$ (2.12 mg, 2.30 µmol, 1.00 eq) and NHS-SS-PEG$_4$-Biotin (3.50 mg, 4.60 µmol, 2.00 eq) were dissolved in 200 µL of DMF. After the addition of triethylamine (0.63 µL, 4.60 µmol, 2.00 eq), the reaction mixture was stirred at room temperature for 1 h. The crude product was purified via preparative reverse phase HPLC. Solvent was removed by lyophilization.

**HPLC** (25% → 75% Buffer B, 70 min, 3.5 ml/min): $R_t$ = 46 min.

**MS** (HR-ESI, pos): measured m/z = 1549.7121, calculated m/z = 1549.7126 for $C_{73}H_{113}N_8O_{22}S_8$ $[M+H]^+$.

**d) Synthesis of TMR-DN**

TMR-DN was synthesized and purified as described in the literature[1]. It is worth mentioning that TMR-DN for imaging applications in cells has to be extremely pure. Since the contact-quenched dyes are non-fluorescent, any fluorescent impurity would dramatically decrease the turn-on ratios and detrimentally affect the live cell imaging experiments. For single molecule experiments, the concentration of the TMR-DN has to be determined precisely. The concentration of the TMR-DN can be calculated by using the extinction coefficients 133,000 $M^{-1}cm^{-1}$ (at 549 nm in trifluoroethanol containing 0.1% TFA) or 118,500 $M^{-1}cm^{-1}$ (at 548 nm in methanol).
